# Mediator Tail Subunits Hierarchically Couple Transcriptional Condensates to Gene Activation and Genome Organization

**DOI:** 10.64898/2026.04.21.719956

**Authors:** Gurranna Male, Suman Mohajan, Linda S. Rubio, David S. Gross

**Author notes:** Equal contribution (Suman Mohajan, Linda S. Rubio).

## Abstract

Cells respond to acute environmental stress by rapidly reorganizing transcriptional machinery and genome architecture, yet how these processes are mechanistically integrated remain poorly understood. We find that Mediator Tail subunits function in a hierarchical fashion to coordinate transcription factor condensate assembly, three-dimensional genome organization and transcriptional output during the heat shock response (HSR) in *Saccharomyces cerevisiae*. We identify the Mediator Tail triad—Med2, Med3, and Med15—as exceptionally enriched in intrinsically disordered regions and possessing strong intrinsic liquid–liquid phase separation potential, in contrast to the more structured Tail subunits Med5 and Med16. Live-cell imaging reveals that this IDR-rich triad is critically required for thermal stress-induced Heat Shock Factor 1 (Hsf1) condensate formation, *HSR* gene coalescence and robust transcriptional induction. Mechanistically, Med15 executes these functions through its activator-binding domains, with the IDR-rich ABD2 playing a dominant role in stabilizing Hsf1, Mediator, and RNA polymerase II (Pol II) occupancy at *HSR* loci, while the C-terminal IDRs of Med2 and Med3 provide critical interaction platforms that couple condensate formation to genome organization. Strikingly, Med16 defines a parallel regulatory axis: although dispensable for Hsf1 condensate nucleation, Med16 is required for Mediator and Pol II condensate formation and for *HSR* gene coalescence, revealing that transcription factor clustering and productive transcriptional condensates are mechanistically separable. Finally, Med5 plays a minor yet detectable role in *HSR* transcription and gene coalescence. Together, our findings establish a modular and hierarchical organization of the Mediator Tail that integrates phase separation, transcriptional condensate composition, and 3D genome architecture to drive rapid stress-induced gene activation.

## Introduction

Mediator is an evolutionarily conserved co-activator of Pol II transcription that plays a central and integrative role in gene expression. Its dynamic and modular architecture, comprising the Head, Middle, and Tail modules (**Fig. 1a**), enables Mediator to interact with gene-specific transcription factors (TFs), Pol II, general transcription factors such as TFIID and TFIIH, and co-activator complexes including SAGA^1–4^. Structural and biochemical studies have demonstrated that Mediator undergoes extensive conformational rearrangements that facilitate high-affinity Pol II engagement and transcriptional activation^5,6^. During pre-initiation complex (PIC) assembly and transcription initiation, Mediator regulates multiple dynamic aspects of transcription, including TF function and multivalent activator interactions, TF residence time on DNA, transcriptional bursting and reinitiation, Pol II distribution across genes, and phosphorylation of the Pol II C-terminal domain (CTD)^1–4^. Beyond its role in initiation, Mediator coordinates post-initiation transcriptional processes, contributing to transcription elongation, termination, and co-transcriptional mRNA splicing and nuclear export^7^ (reviewed in ^3,4,8^). In addition to these transcriptional functions, Mediator interfaces with chromatin regulatory pathways, influencing histone and DNA methylation and contributing to epigenetic transcriptional memory^3,4,9^.

**Figure 1.**
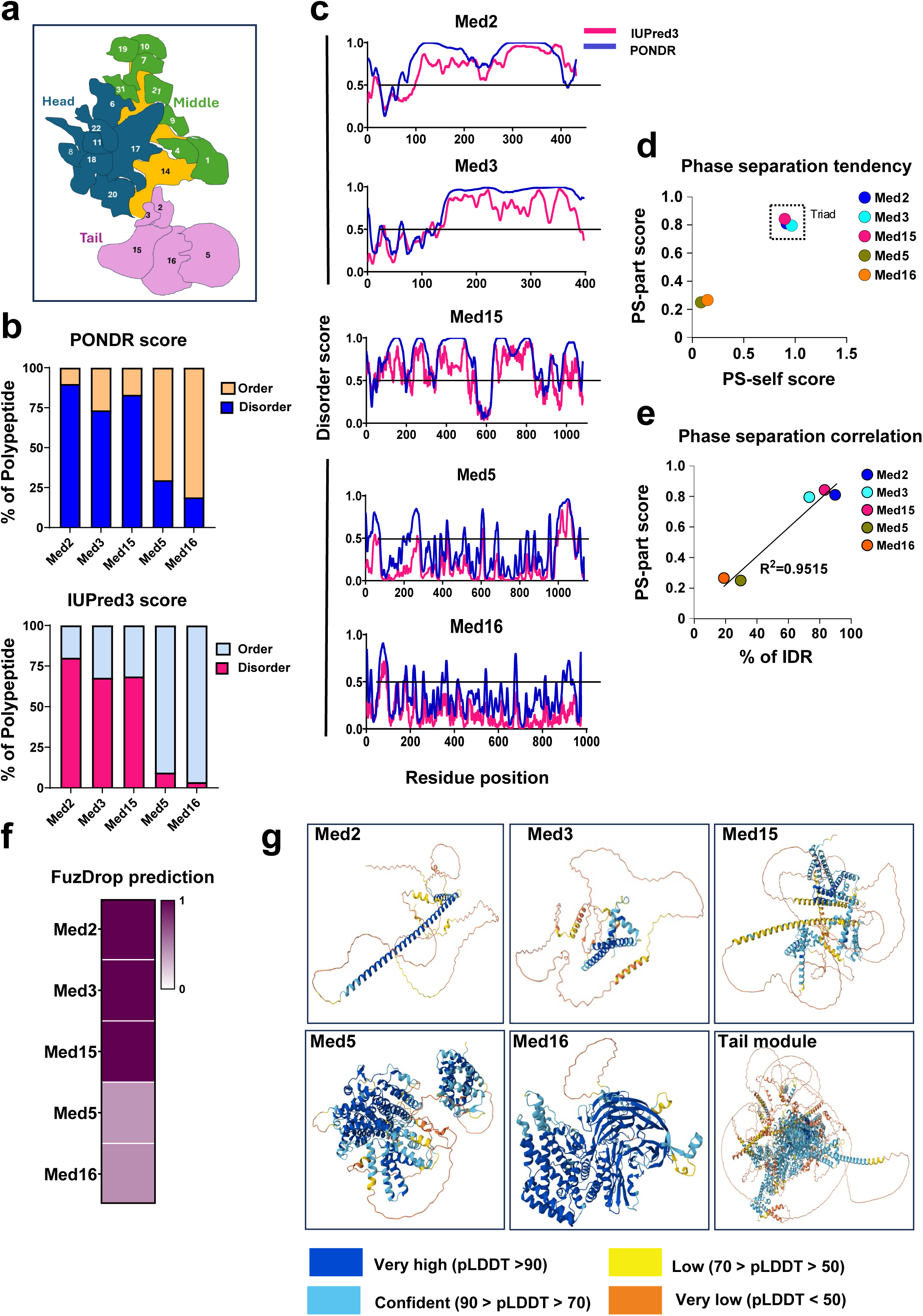
| Mediator Tail subunits Med2, Med3, and Med15 exhibit high intrinsic disorder and a strong predicted propensity for liquid–liquid phase separation. **a**, Schematic of the *Saccharomyces cerevisiae* Mediator complex illustrating its modular organization and subunit composition (adapted from Philip *et al.*, *eLife*, 2015 (ref.5). **b,** Intrinsic disorder predictions for the indicated Mediator Tail subunits based on PONDR and IUPred3 analyses; residues with prediction scores ≥0.5 are classified as disordered. **c,** Comparative disorder prediction profiles of the indicated Mediator Tail subunits generated using IUPred3 (magenta) and PONDR VSL2 (blue). **d,** Phase separation propensity of Mediator Tail subunits predicted by PhaSePred, shown as PS-part and PS-self scores; the Mediator Tail subunit triad (Med2, Med3, and Med15) is highlighted (dotted box). **e,** Correlation between PS-part scores (PhaSePred) and the proportion of intrinsically disordered residues (PONDR) across the indicated Mediator subunits. **f,** Heatmap of predicted liquid–liquid phase separation probabilities for Mediator Tail subunits calculated using FuzDrop, with darker shading indicating higher propensity. **g,** Predicted structures of individual Mediator Tail subunits and the assembled Tail module generated using AlphaFold3. pLDDT, Predicted Local Distance Difference Test.

Beyond transcription, Mediator plays a pivotal regulatory role in genome organization by orchestrating short– and long-range chromatin interactions through partnerships with diverse factors, including chromatin architectural proteins and remodelers^4,8^. By interacting with cohesin, Mediator promotes enhancer-promoter interactions and higher-order chromatin folding to regulate global chromatin structure^10–12^. These observations, primarily made in mammalian systems, reflect developmental processes. Less well understood is the role Mediator might play in the dynamic 3D genome restructuring that occurs in response to environmental stress^13–17^.

The robust heat shock response (HSR) system in budding yeast provides a powerful model to uncover the basic mechanisms of genome organization and gene regulation. The HSR is a conserved transcriptional program that safeguards eukaryotic cells against proteotoxic stress and is orchestrated by the transcription factor, Heat Shock Factor 1 (Hsf1)^18,19^. Upon activation by stressors such as heat, ethanol, or oxidative stress, Hsf1 undergoes trimerization, binds to heat shock elements (HSEs) upstream of *HSR* genes, and induces expression of molecular chaperones and cytoprotective proteins^18,20–22^. This regulatory mechanism is evolutionarily conserved, as evidenced by the functional interchangeability of human HSF1 and its yeast counterpart^23^.

Reversible formation of biomolecular condensates, a process that frequently involves liquid-liquid phase separation (LLPS), has emerged as a key mechanism regulating the cellular stress response from yeast to mammals^15,16,24^ (reviewed in ^19,25^). This process is primarily driven by intrinsically disordered regions (IDRs). Notably, Mediator contains extensive IDRs in many subunits, consistent with its striking conformational flexibility ^26,27^. Previous work has suggested that dynamic IDR-IDR interactions between TFs and Mediator can drive the formation of phase-separated condensates^28,29^, yet roles played by such IDR-driven interactions in fostering recruitment of regulatory factors, gene-specific transcription, and 3D genome organization remain unclear.

Recent studies have shown that Hsf1 forms reversible nuclear condensates in response to acute thermal stress and that these localize to *HSR* gene loci in both human and yeast ^16,24^. In *Saccharomyces cerevisiae*, these heat-inducible transcriptional condensates drive *HSR*-*HSR* gene interactions across and between chromosomes^14–16^. In addition to driving profound reorganization of genome structure, such condensates concentrate components of the transcriptional machinery, including Hsf1, Pol II, and Mediator^16,31,33^. However, whether factors beyond Hsf1 drive the formation of transcriptional condensates and concomitant 3D genome reorganization, and the mechanistic basis by which they might do so, are largely unknown.

Using the HSR system in budding yeast, we previously showed that the Mediator Tail triad, comprised of Med2-Med3-Med15, can be independently recruited to *HSR* genes through interactions with Hsf1, which uses its dual activation domains to physically interact with Med15 ^34,35^. However, the role of Mediator in heat shock-induced transcriptional condensate formation and *HSR* gene clustering has remained unclear. Here, we demonstrate that subunits within the Tail triad are critically required for heat shock-induced formation of transcriptional condensates, *HSR* gene activation, and concomitant 3D genome restructuring, and principally act through their intrinsically disordered regions. A fourth, highly structured subunit, Med16, defines a parallel regulatory axis: although dispensable for Hsf1 condensate nucleation, Med16 is required for Mediator and Pol II condensate formation and for *HSR* gene coalescence, revealing that transcription factor clustering and productive transcriptional condensates are mechanistically separable. Med5, another highly structured Tail subunit, plays comparatively minor roles in stress-induced genome restructuring and *HSR* gene expression. Our findings establish a modular and hierarchical organization of the Mediator Tail that integrates phase separation, transcriptional condensate composition, and 3D genome architecture to drive rapid stress-induced gene activation.

## Results

### Mediator Tail triad subunits exhibit robust intrinsic LLPS potential

The formation of biomolecular condensates is often driven by LLPS and is dictated by IDR abundance, amino acid composition, and sequence patterning^36–39^. We therefore systematically evaluated Mediator Tail subunits for IDR content, underlying amino acid composition, and intrinsic LLPS potential. IDR abundance and LLPS propensity were quantified using complementary sequence-based and machine-learning approaches for predicting intrinsic disorder and droplet-forming potential^40–44^. This analysis revealed that Med2, Med3, and Med15 exhibit a high degree of disorder, with >70% of each polypeptide predicted to be disordered **(Fig. 1b**). Med2 and Med3 contain extended IDRs, whereas Med15 features multiple IDRs that punctuate its sequence end-to-end **(Fig. 1c)**. In contrast, Med5 and Med16 contain few regions predicted to be disordered.

To evaluate the phase-separation potential of Mediator Tail subunits, we used PhasePred to predict both self-assembling (PS-Self) and partner-dependent (PS-Part) propensities. Med2, Med3, and Med15 emerged as dominant candidates, exhibiting markedly elevated PS-Self and PS-Part scores and the highest predicted phase-separation propensity among all Mediator subunits (**Figs. 1d and S1 a, b**). Notably, PS-Part scores strongly correlated with IDR content (R² = 0.9515), implicating IDRs within the Tail triad as major determinants of phase separation (**Fig. 1e**). Consistent with these findings, FuzDrop analysis also predicted elevated LLPS propensity for Med2, Med3, and Med15 (**Fig. 1f**). AlphaFold3 structural predictions^45^ likewise suggested extensive unstructured regions in these subunits, in contrast to the more ordered architectures of Med5 and Med16 (**Fig. 1g**). Together, these data indicate that the Mediator Tail triad is characterized by pronounced intrinsic disorder and a strong propensity for phase separation.

Because amino acid composition is a critical determinant of IDR-driven phase separation^37^, we next analyzed the residue composition of Mediator Tail subunits. The Tail triad was markedly enriched in asparagine (N) and glutamine (Q) residues, with Med2 showing preferential enrichment for asparagine, Med15 for glutamine, and Med3 for both. By contrast, Med5 and Med16 exhibited minimal enrichment in these residues (**Fig. S1c, d**). Notably, N and Q residues were largely spatially aligned with IDRs within the Tail triad (**Fig. S1e**), and their combined abundance was significantly higher than that observed in other Tail subunits. Collectively, these features indicate that the Mediator Tail triad harbors intrinsic sequence and structural properties consistent with robust liquid–liquid phase separation potential.

### The Mediator Tail triad regulates Hsf1 condensate formation in response to heat shock

HSF1, the master regulator of the heat shock response, undergoes LLPS to form nuclear condensates at *HSR* gene loci in both yeast and human cells^16,24^ and in yeast drives the 3D reorganization of *HSR* genes^15,16,31,46^. Recently, evidence has been reported for HSF1-mediated 3D reorganization of *HSR* genes in mouse embryonic fibroblasts^47^. Although Mediator and Pol II colocalize with Hsf1 following exposure to acute thermal stress^16,31,33^, the contribution of individual Mediator subunits remains poorly defined. To investigate the role of the Mediator Tail in the HSR, we generated haploid yeast strains harboring individual gene deletions (*med2Δ, med3Δ*, *med15Δ*, *med5Δ*, *med16Δ*) and assessed growth phenotypes at 24°, 30°, and 37°C. Except for *med5Δ*, all mutants exhibited sensitivity to the elevated temperature (37°C) (**Fig. S2**), implicating essential roles for Med2, Med3, Med15, and Med16 in the HSR.

Dynamic IDR-IDR interactions of transcriptional machinery drive phase-separated condensates, crucial for transcriptional regulation (reviewed in ^36,48,49^). Optimal levels of TF IDR-IDR interactions facilitate condensate formation that often correlates with transcriptional induction^28,29,48–50^ (although see ref.^51^). Mammalian Med1 plays an essential role in condensate formation in mammalian cells, facilitating the compartmentalization of Pol II and positive regulators and activating transcription by concentrating transcriptional machinery via its large IDR^28,29,39,52^. To test the role of Mediator Tail subunits in HS-induced Hsf1 condensate formation, we employed live-cell imaging of wild-type (WT) and individual Tail subunit deletion strains expressing Hsf1-mNeonGreen (Hsf1-mNG). Consistent with previous observations^15,16,31,33,46^, Hsf1-mNG rapidly condensed in WT cells into several discrete intranuclear puncta in response to a 2.5-minute, 25° to 39°C thermal upshift. Such puncta were dynamic and began to resolve as early as 15 min (**Fig. 2a**).

**Figure 2.**
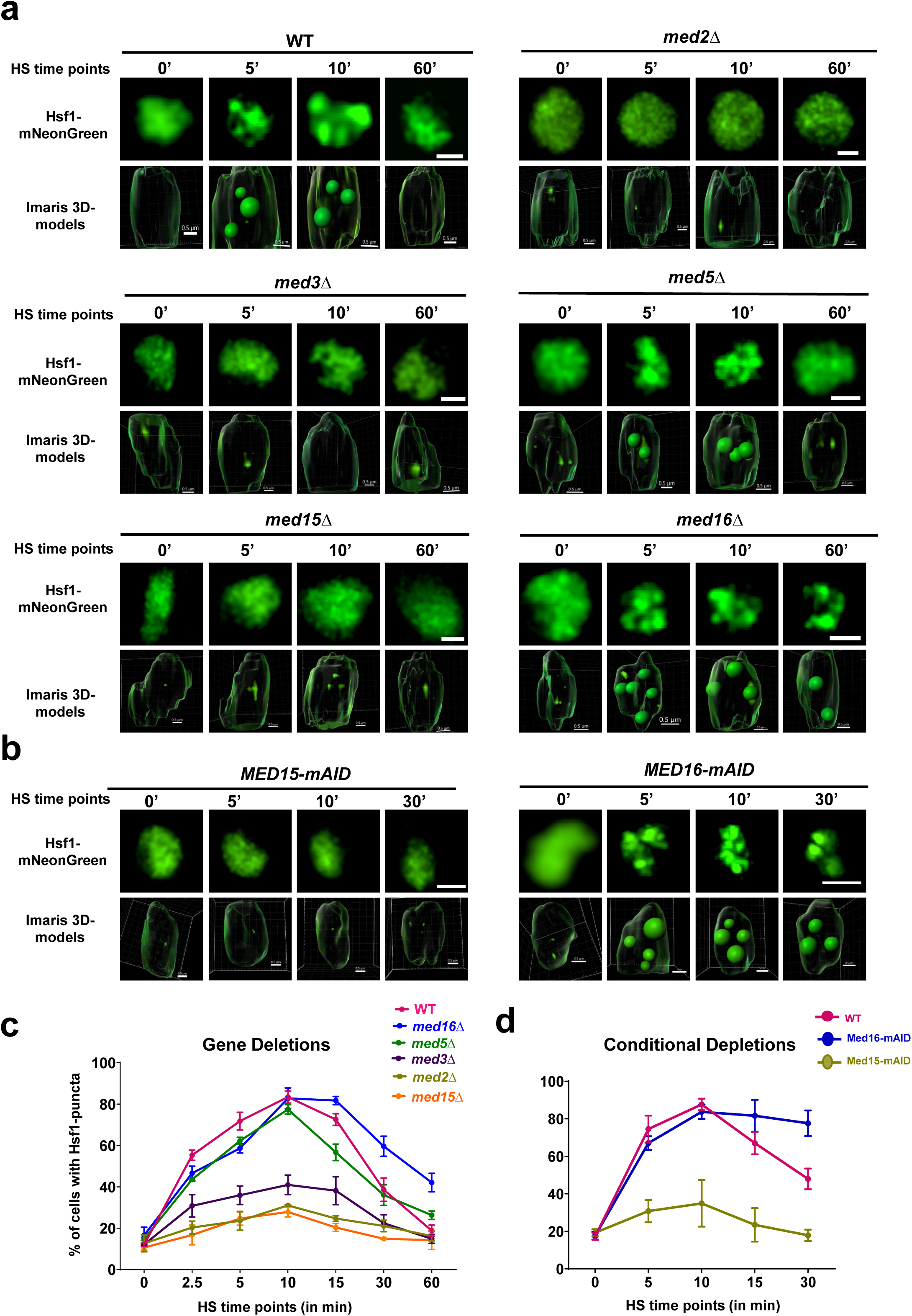
| Mediator Tail subunits regulate Hsf1 condensate kinetics during heat shock. **a**, Representative fluorescence micrographs and corresponding three-dimensional reconstructions (Imaris v10.2.0, Cells module) of Hsf1 condensates in the indicated wild-type (WT) and deletion strains during a 39°C heat shock time course. 0 min, cells maintained at 25°C. Scale bars, 1 µm; for Med15-mAID and Med16-mAID, 2 µm. Scale bar for Imaris-rendered 3D models, 0.5 µm. **b,** As in **a**, except cells expressing the indicated mAID-tagged proteins were imaged. Prior to initiating heat shock, cells were treated with 1 mM IAA for 60 min at 25°C. Scale bars, 2 µm. **c,d,** Quantification of the fraction of cells containing two or more Hsf1 condensates (n ≥ 2) during heat shock. Data represent the mean of two independent biological replicates; error bars indicate s.d.

Deletion of any component of the Mediator Tail triad markedly suppressed Hsf1 condensate formation, as reflected by both the number of puncta per cell and the fraction of cells containing puncta, although the kinetics of formation and resolution were similar to those observed in WT cells (**Figs. 2a,c; S3i)**. In contrast, deletion of *MED5* or *MED16* did not reduce either the kinetics or abundance of intranuclear Hsf1 condensates. Notably, Hsf1-mNG condensates persisted longer in *med16Δ* cells than in WT cells (**Fig. 2c**). Consistent with these findings, conditional depletion of Med16 using the auxin-inducible degron (mAID) system did not suppress Hsf1 clustering but instead prolonged condensate persistence, whereas AID-mediated Med15 depletion abolished Hsf1 condensation (**Figs. 2b,d; S3h,j**). Auxin treatment efficiently reduced Med15 and Med16 protein levels (∼99% and ∼90%, respectively) within 30 minutes, without affecting cell viability over the time course analyzed (**Fig. S3a–f**).

Together, these results demonstrate that the Mediator Tail triad, which is enriched in IDRs and exhibits strong LLPS propensity, contributes to the HSR by stimulating the formation of Hsf1 condensates. By contrast, Med5 and Med16, both of which lack pronounced IDRs and LLPS propensity (**Fig. 1**), are dispensable for Hsf1 nucleation.

### Both IDR-containing and structured Mediator Tail subunits drive HS-induced *HSR* gene interactions

Heat shock–induced transcriptional condensates containing Hsf1, Pol II, and Mediator are proposed to drive the spatial reorganization of *HSR* genes, with evidence supporting a functional link between Hsf1 condensate formation and intergenic *HSR* gene interactions^16,31^. To gain further insight into the relationship between condensate assembly and *HSR* gene coalescence, we analyzed the contribution of Mediator Tail subunits using a highly sensitive version of chromosome conformation capture, TaqI-3C ^53^, in both gene deletion and conditional depletion strains (strategy summarized in **Fig. 3a**). This analysis included the IDR-rich subunits Med2, Med3, and Med15, as well as the more structured subunits Med5 and Med16. In WT cells, brief heat shock (5 min) induced robust *cis-* and *trans*-intergenic interactions among *HSR* genes (**Figs. 3b** and **S4**). Deletion of *MED2, MED3* or *MED15*, each of which disrupts Hsf1 condensate formation, resulted in a pronounced loss of *HSR* intergenic contacts. Notably, deletion of *MED16* caused the most severe defect in *HSR* gene coalescence despite leaving Hsf1 condensates intact. Deletion of *MED5* had a more variable and less impactful effect on both *cis-* and *trans*-interactions (**Figs. 3b** and **S4**).

**Figure 3.**
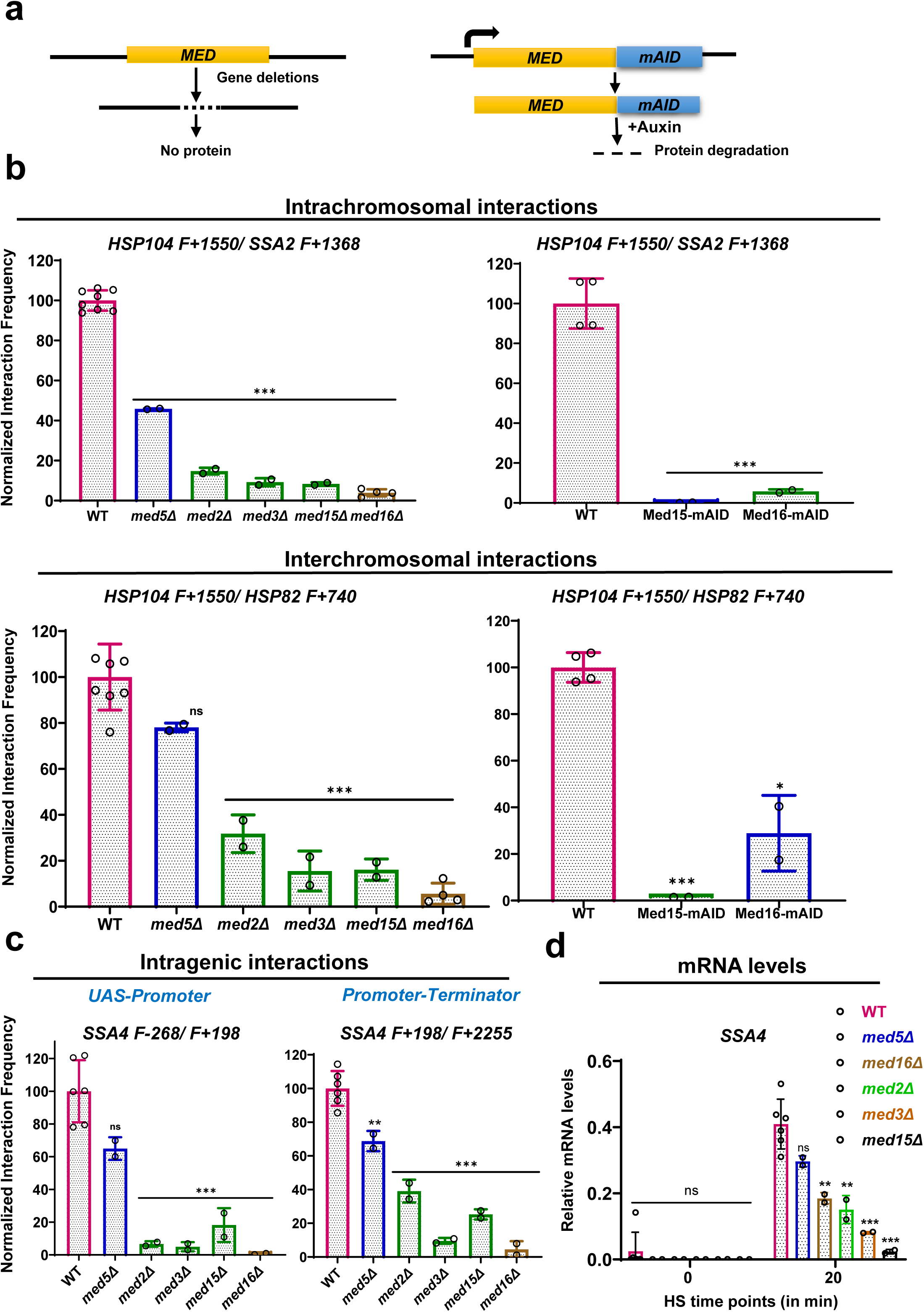
| Mediator Tail subunits drive heat-shock-induced *HSR* gene interactions and promote *HSR* mRNA abundance. **a**, Schematic overview of the orthogonal experimental strategies—gene deletion and conditional depletion—used to assess the role of Mediator Tail subunits in *Heat Shock Response* (*HSR*) gene interactions. **b, c,** Normalized interaction frequencies of *HSR* genes in the indicated strains subjected to a 5 min heat shock at 39°C. **b** shows representative cis– and trans-intergenic interactions, and **c** shows intragenic interactions. Pairwise tests used forward (F; sense-strand) primers positioned nearby the indicated Taq I site; primer locations are illustrated in Fig. S1 of Mohajan et al (ref. 33). Data are presented as mean ± s.d.; N = 2-8 independent samples (as indicated). Statistical significance relative to WT was determined using an unpaired, two-tailed *t*-test (*, *P* < 0.05; **, *P* < 0.01; ***, *P* < 0.001; ns, not significant). Location of gene loci: *HSP104* and *SSA2*, Chr XII; *HSP82*, Chr XVI; *HSP12*, Chr VI; *SSA4*, Chr V. **d,** Relative *SSA4* mRNA abundance measured by RT–qPCR. Data represent mean ± s.d. from the indicated number of independent biological replicates. Statistical significance relative to WT was assessed as above.

To validate the most striking phenotypes, we conditionally depleted Med15 or Med16 using AID as above. Like *med15Δ*, conditional depletion of Med15 severely disrupted both *cis-* and *trans-HSR* intergenic interactions (**Figs. 3b, S5a**). Med16-mAID produced a similar but weaker defect than *med16Δ* that may reflect the incomplete depletion of Med16 protein (**Figs. 3b, S3d-e, S5b**).

We next examined intra-locus *HSR* interactions, including enhancer–promoter (E-P), promoter–coding region, and promoter–terminator (gene-looping) contacts. Except for Med5, all Mediator Tail subunits were required for efficient intragenic interactions in acute HS cells (**Figs. 3c, S4c**). Conditional depletion of Med16 yielded comparable results to *med16Δ* (**Fig. S5b**). The similar impact of gene deletion and conditional protein depletion of Med15 and Med16 argues against the effects observed having arisen from either extragenic suppression or auxin-induced effects. Together, these orthologous approaches argue for a dual role for IDR-containing Med15 in promoting Hsf1 clustering and *HSR* intergenic interactions and a singular role for Med16 in promoting HS-induced 3D genome restructuring.

### Mediator Tail triad subunits drive robust *HSR* gene transcription

TF condensates have been proposed as a key mechanism for transcriptional regulation^28,29,32,54^. In yeast, *HSR* gene transcription is temporally correlated with HS-induced Hsf1 condensation^15,16,31,33,46^. To determine whether Mediator Tail subunits implicated in Hsf1-condensate formation and *HSR* gene coalescence are required for HS-induced *HSR* transcription, we measured mRNA levels of representative *HSR* genes (*HSP104, SSA4,* and *BTN2*) during a HS time course using RT–qPCR. Loss of Tail triad subunits Med2, Med3, and Med15 severely impaired *HSR* gene transcription, with Med15 exhibiting the most pronounced defect (**Figs. 3d, S6**). These transcriptional defects parallel the disruptions observed in Hsf1 condensate formation and *HSR* gene interactions. In contrast, deletion of Med16, which strongly disrupts *HSR* gene interactions but not Hsf1 condensates, caused only a moderate reduction in *HSR* transcription. Finally, deletion of Med5, which does not affect the number of Hsf1 condensates per cell and has a variable impact on *HSR* intergenic interactions, had a modest yet detectable effect on *HSR* gene transcription. Together, these data indicate that the Tail triad subunits are essential for robust *HSR* transcription, that Med16 plays a moderate but supportive role, and that Med5 plays a minor role.

### Med15 activator-binding domains govern the HSR through regulation of Hsf1, Mediator, and Pol II occupancy

Med15 is a well-characterized Mediator subunit that regulates gene expression through interactions with transcription factors across eukaryotes, from yeast to human^34,55–58^. Yeast Med15 contains three N-terminal activator-binding domains (ABDs 1-3) comprised of extensive regions predicted to be intrinsically disordered (**Figs. 4a, S7**) and that are essential for TF binding ^56,59,60^. Through these domains, Med15 engages TFs via a “fuzzy” protein interface^57,58,60,61^. Med15 additionally harbors an evolutionarily conserved, N-terminal TF-interacting KIX domain^55,56^ and a C-terminal Mediator-associated domain (MAD) that mediates interactions with the remainder of the Mediator complex, TFIIE and TFIIH^5,62^. As a core component of the Mediator Tail, Med15 has been shown to be essential for the HSR^34,35^, a notion confirmed and extended by observations described above (**Figs. 2, 3, S4-S6**).

**Figure 4.**
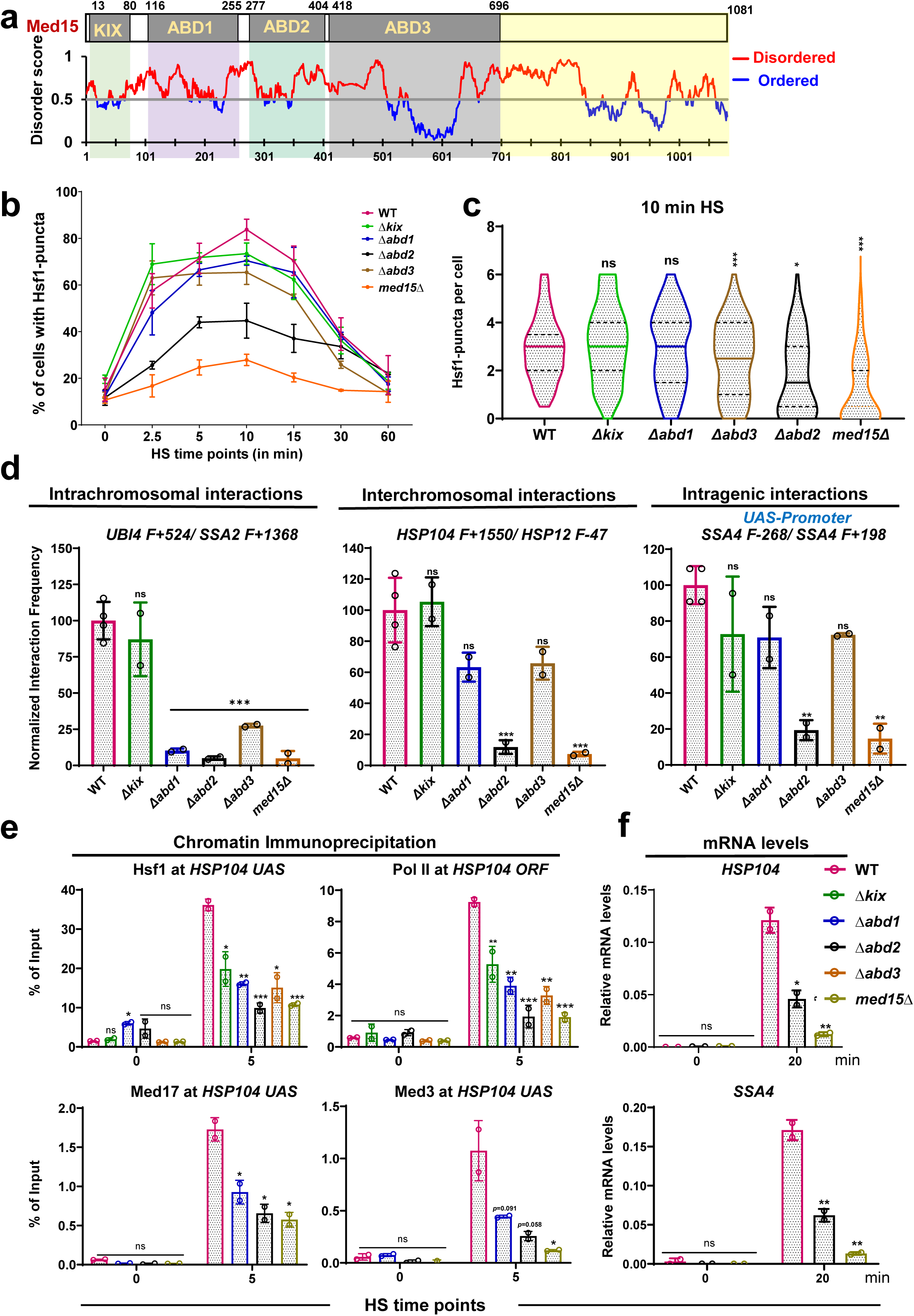
| The Med15 ABD2 domain is required for Hsf1 condensate formation and heat-shock-induced *HSR* gene interactions. **a**, Schematic of Med15 domain architecture and deletion constructs, with the domains deleted indicated. Predicted intrinsic disorder across Med15 was assessed by IUPred, with amino acid positions shown. **b,** Quantification of cells containing two or more Hsf1 condensates in WT and Med15 domain deletion strains during heat shock using Imaris v10.1.0 (Cells module). Data represent the mean of two independent biological replicates; error bars indicate s.d. **c,** Violin plot showing the distribution of Hsf1 puncta per cell in WT and *med15* mutants after 10 min of heat shock, quantified as in **b**. Horizontal lines indicate median values; dashed lines indicate quartiles. Statistical significance relative to WT was assessed by one-way ANOVA (*, *P* < 0.05; **, *P* < 0.01; ***, *P* < 0.001). **d,** Normalized interaction frequencies of *Heat Shock Response* (*HSR*) genes measured by Taq I – 3C in the indicated strains after 5 min HS, showing intrachromosomal (cis), interchromosomal (trans), and intergenic interactions. Data represent mean ± s.d. from two independent biological replicates. Statistical significance relative to WT was determined using unpaired two-tailed *t*-test (*, *P* < 0.05; **, *P* < 0.01; ***, *P* < 0.001). **e,** ChIP analysis of Hsf1, Pol II large subunit Rpb1, and Mediator subunits Med17 (Head) and Med3 (Tail) at the *HSP104* locus in the indicated strains. Data represent mean ± s.d. from two independent biological replicates. Statistical significance relative to WT was assessed as in **d**. **f,** Relative mRNA levels of *HSP104* and *SSA4* measured by RT–qPCR at the indicated heat-shock time points in the indicated strains. Data represent mean ± s.d. from two independent biological replicates.

To determine whether Med15 mediates these HSR functions through its ABDs, we generated haploid strains carrying individual genomic deletions of each domain^59^ (see Methods). Deletion of ABD2 (*Δabd2*, residues 277–404) caused an increasingly severe growth defect as temperature was elevated, although not as severe as that conferred by *med15Δ* (**Fig. S8a-c**), whereas the remaining ABD deletions had little detectable effect. Significantly, none of the domain deletions altered overall Med15 protein expression levels (**Fig. S8d**).

We next assessed whether the ABD deletions affected Hsf1 condensate formation. Except for *Δabd2*, none of the deletions significantly altered the fraction of cells with Hsf1 or kinetics of formation over the heat shock time course (**Fig. 4b**). Quantification of Hsf1 puncta per cell revealed a substantial reduction in the *Δabd2* mutant, a moderate reduction in *Δabd3*, and no appreciable effect in either the *Δkix* or *Δabd1* strains (**Fig. 4c**). We then examined the contribution of Med15 ABDs to intergenic interactions at *HSR* genes by 3C. All ABD deletions impaired intrachromosomal (*cis*) interactions, whereas only *Δabd2* consistently disrupted interchromosomal (*trans*) interactions and intragenic contacts (**Figs. 4d and S9a**). Moreover, analysis of intragenic architecture showed that ABD2 is required for both enhancer–promoter (E–P) and promoter–terminator (gene-looping) interactions. In contrast, deletion of the KIX domain, ABD1, or ABD3 resulted in only moderate reductions in gene-looping interactions, while leaving E–P interactions largely intact. Together, these findings indicate that all Med15 ABDs contribute to the 3D genome topology of *HSR* genes, with ABD2 playing the predominant role.

To determine how Med15 contributes to the HSR through its ABDs, we assessed the occupancy of key HSR transcriptional components, including Hsf1, Pol II (Rpb1), and the Mediator Head (Med17) and Tail (Med3) subunits, using ChIP at representative *HSR* gene loci. All Med15 ABDs supported recruitment of these factors, with ABD2 typically playing the most important role (**Figs. 4e, S10**). It is notable that Med15, acting through each of its activator binding domains, enhances HS-induced Hsf1 recruitment in addition to that of Pol II and Mediator. This observation is consistent with previous reports suggesting that Med15 can facilitate (or stabilize) the binding of gene-specific transcription factors to their cognate sites in the yeast genome^34,63^. We further evaluated the role of ABD2 in driving *HSR* gene expression and found that its deletion severely reduced mRNA abundance of *HSR* genes (**Figs. 4f, S9b**). Together, these results indicate that Med15 ABDs are critical for the HSR, functioning to enhance or stabilize transcriptional machinery at *HSR* genes in an HS-dependent manner, with ABD2 playing the most important role.

### The C-terminal IDRs of Med2 and Med3 regulate the HSR by modulating HSF1 condensation, genome topology and factor occupancy

Previous studies have demonstrated that Hsf1 is required for the HS-induced formation of transcriptional condensates, 3D genome restructuring, and *HSR* gene activation in acutely stressed cells^15,16,31,46^. Consistent with this framework, our results show that each component of the Mediator Tail triad—Med2, Med3, and Med15—is essential for a robust HSR, as loss of any individual subunit severely compromises the response. These findings support a model in which the Tail triad subunits function cooperatively as an integrated triad rather than as independent factors^5,64–66^. Furthermore, Med15 contributes to the HSR through its activator-binding domains (ABDs), with ABD2 playing a predominant role (**Figs. 4, S9, S10**). To define the IDR-mediated interactions that underlie Tail triad assembly, we generated intermolecular interaction maps using the FINCHES online tool^67^. These analyses predicted strong interactions between Med15 ABDs 1–3 and the C-terminal IDRs of Med2 (residues 290–400) and Med3 (residues 301–374) (**Fig. S11**). In contrast, the same ABDs showed only weak to moderate predicted interactions with Med5 or Med16.

To determine whether Med2 and Med3 contribute to the HSR through their C-terminal IDRs, we used CRISPR/Cas9 to generate genomic deletions of these glutamine– and asparagine-enriched regions (see **Fig. 5a**). The *med2-ΔIDR* mutant grew poorly at elevated temperature while *med3-ΔIDR* evinced no growth defect (**Fig. S12a**). Notably, deletion of the C-terminal IDRs did not alter the protein expression levels of either *med2-ΔIDR* or *med3-ΔIDR* (**Fig. S12b, c**).

**Figure 5.**
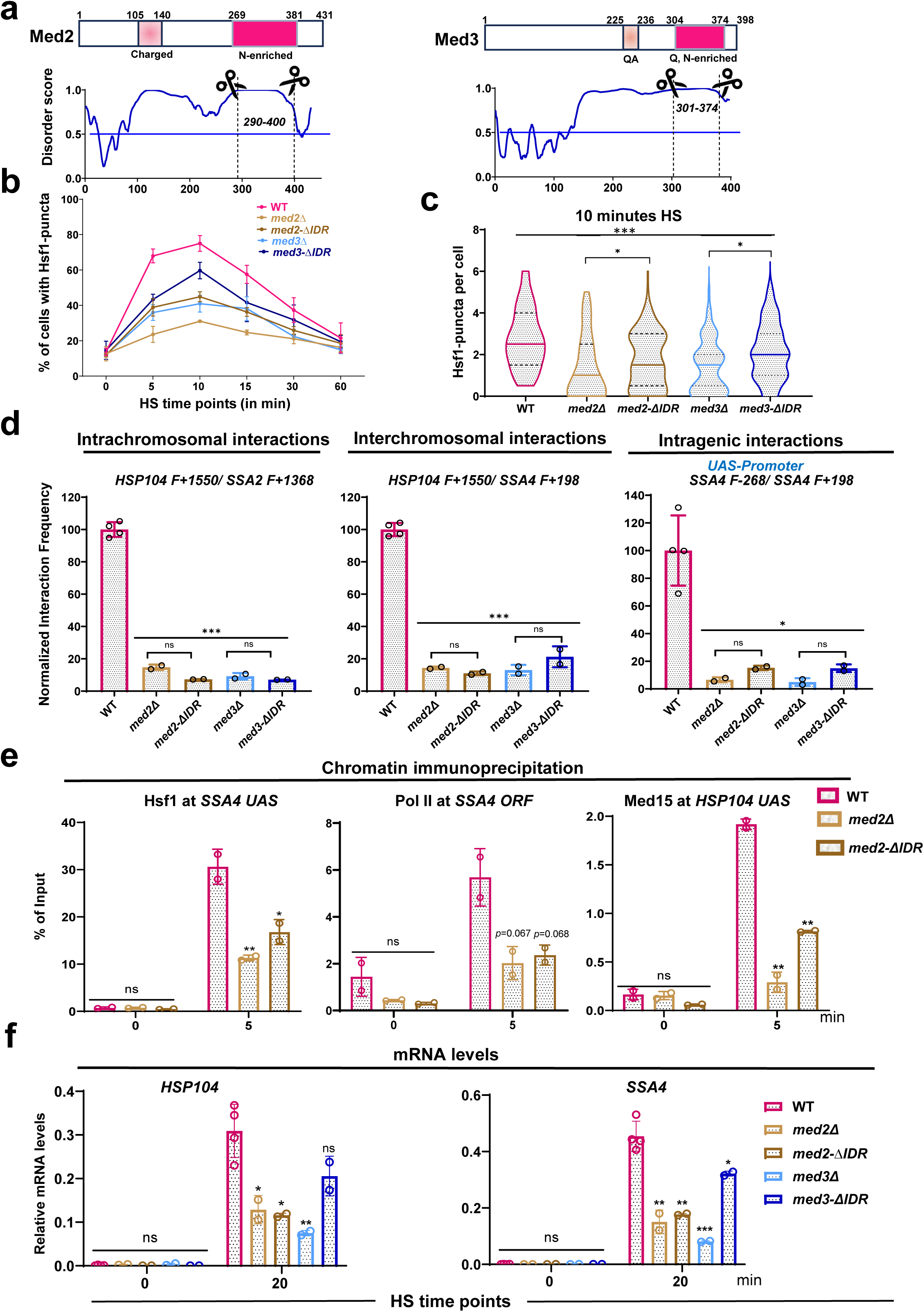
| Med2 and Med3 C-terminal IDRs regulate the heat-shock response by modulating Hsf1 clustering, genome topology, and Hsf1, Mediator and Pol II occupancy. **a**, Domain architecture of Med2 and Med3 with accompanying IUPred3 analysis. Dashed lines indicate locations of CRISPR-Cas9–targeted deletions of the C-terminal IDRs. **b,** Hsf1 puncta formation in WT and Med2 or Med3 gene / domain deletion strains during a heat shock time course. **c,** Violin plot showing the distribution of Hsf1 puncta per cell after a 10 min heat shock, quantified using Imaris. Solid horizontal lines indicate median values. Discontinuous lines depict quartiles. Statistical significance relative to WT was determined by one-way ANOVA. *, *P* < 0.05; **, *P* < 0.01; \*\*\**P* < 0.001. **d,** Normalized interaction frequencies of *Heat Shock Response* (*HSR*) genes measured by Taq I – 3C in the indicated strains after 5 min HS, showing intrachromosomal (cis), interchromosomal (trans), and intergenic interactions. Data represent mean ± s.d. (*n* = 2). Statistical significance relative to WT was determined using an unpaired two-tailed *t*-test (*, *P* < 0.05; **, *P* < 0.01; \*\*\**P* < 0.001). **e,** ChIP analysis of Hsf1, Rpb1 and Med15 at representative *HSR* genes in the indicated strains. Data represent mean ± s.d. (*n* = 2). **f,** Relative *HSP104* and *SSA4* mRNA levels measured by RT–qPCR in the indicated strains following heat shock. Data represent mean ± s.d. from independent biological replicates (*n*=2). Statistical significance relative to WT was determined using an unpaired two-tailed *t*-test (*, *P* < 0.05; **, *P* < 0.01; \*\*\**P* < 0.001).

We next examined the impact of these deletions on Hsf1 condensate dynamics during a heat shock time course. Both *med2-ΔIDR* and *med3-ΔIDR* mutants exhibited defects in Hsf1 condensate formation throughout the HS time course, paralleling the more pronounced phenotypes of their corresponding gene deletions (**Fig. 5b**). At 10 min HS, both *med2-ΔIDR* and *med3-ΔIDR* mutants displayed significant reductions in the number of Hsf1 condensates relative to WT. However, the defects were less pronounced than those observed in the corresponding gene deletion strains (**Fig. 5c**).

Next, we assessed the role of the Med2 and Med3 C-terminal IDRs in driving *HSR* gene interactions using the 3C method at the 5-min HS time point. Both *med2-ΔIDR* and *med3-ΔIDR* exhibited severe defects in *HSR* gene coalescence, disrupting *cis-, trans-,* and intragenic interactions (**Figs. 5d** and **S13a**). These defects closely mirrored the phenotypes of the gene deletion mutants, indicating that Med2 and Med3 play a vital role in driving *HSR* gene coalescence through their C-terminal IDRs.

To further define how the Med2 C-terminal IDR influences Hsf1 condensate formation and HSR gene interactions, we assessed the occupancy of Hsf1, Pol II (Rpb1), and Med15 at representative *HSR* genes under NHS and HS conditions. Deletion of *MED2* markedly reduced recruitment of all three factors, and the *med2-ΔIDR* produced a comparable reduction of Rpb1 and Med15 at each locus examined (**Figs. 5e, S13b–d**). We next assessed the contribution of the Med2 and Med3 C-terminal IDRs to HSR gene transcription during the heat shock time course by RT–qPCR. Deletion of the Med2 C-terminal IDR (*med2-ΔIDR*) markedly impaired induction of *HSR* transcripts, including *HSP104* and *SSA4*, phenocopying the *MED2* deletion, whereas *med3-ΔIDR* caused a more modest transcriptional defect relative to *med3Δ* (**Figs. 5f, S14**). Together, these results suggest that Med2 and Med3 regulate the HSR through their C-terminal IDRs, with Med2 playing a dominant role in coordinating Hsf1 condensation, *HSR* gene coalescence, and transcriptional activation.

### Med16 Is required for the formation of functional transcriptional condensates

Observations described above support a model in which Tail triad–mediated multivalent interactions integrate condensate assembly with higher-order chromatin organization to drive robust transcription. By contrast, *med16Δ* profoundly disrupted *HSR* gene coalescence without affecting Hsf1 condensate formation **(Figs. 2, 3, S4)**, consistent with the notion that genome reorganization occurs downstream of TF clustering^16,46^. To determine how Hsf1 condensates in the *med16Δ* mutant differ from those in WT cells, we performed live cell colocalization analyses to define the contribution of individual Mediator subunits to transcriptional condensate composition. Heat shock-induced WT cells exhibited robust recruitment of Mediator and Pol II into Hsf1 condensates, resulting in dynamic transcriptional assemblies. Deletion of *MED5* had little effect on condensate formation, whereas loss of either *MED2* or *MED3* severely disrupted Hsf1, Mediator, and Pol II condensates (**Fig. 6a-d, g-j**). Although *med5Δ* did not alter the fraction of cells containing Hsf1, Mediator (Med15), or Pol II (Rpb3) condensates, it modestly reduced the number of Mediator puncta per cell (**Fig. 6d**).

**Figure 6.**
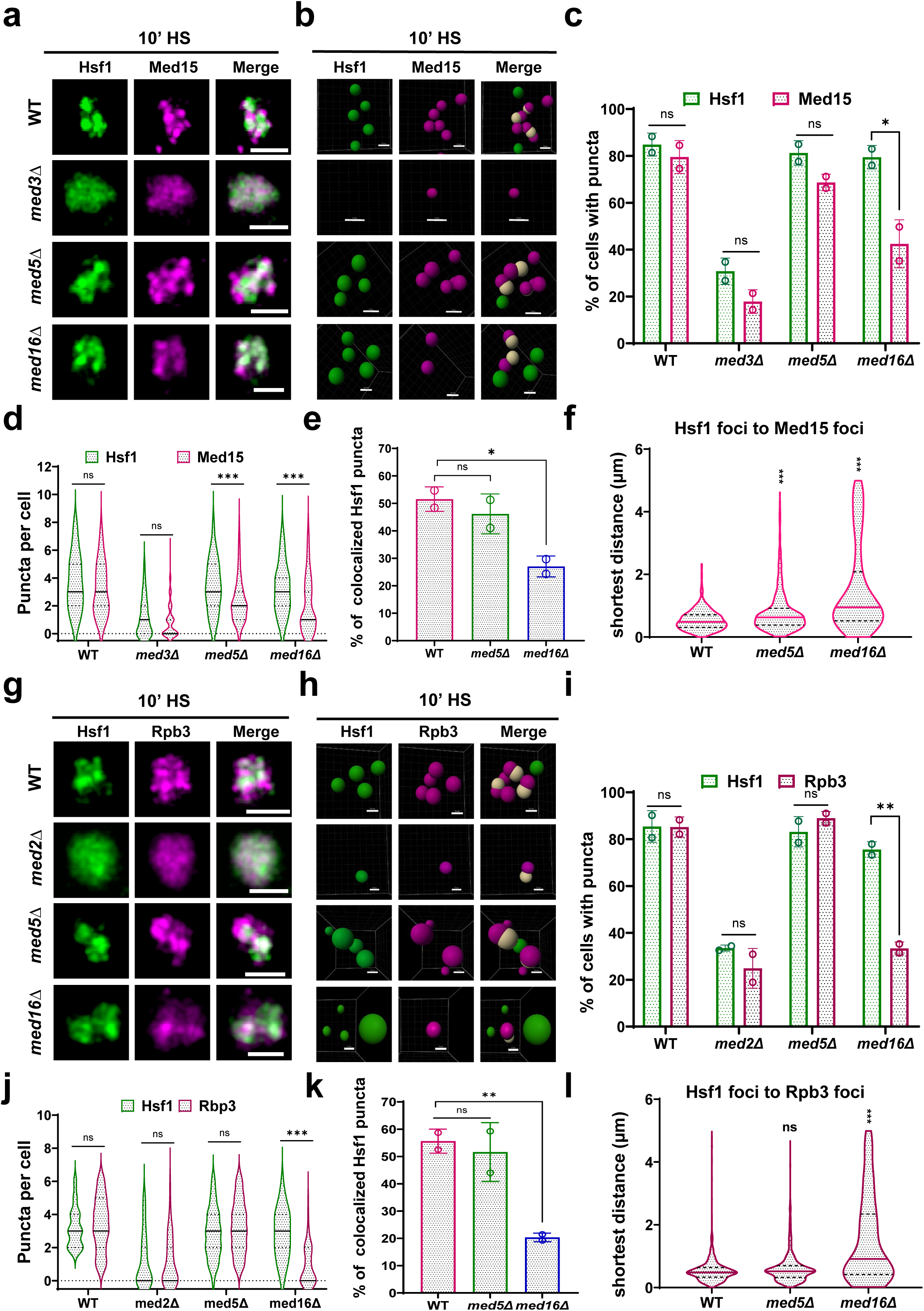
| Med16 is required for HS-induced formation of Mediator and Pol II condensates. **a, g**, Representative live-cell fluorescence images of the indicated strains expressing Hsf1–mNG, Med15–mCherry and Rpb3–mCherry, marking Hsf1, Mediator, and Pol II, respectively, after 10 min HS. Scale bars, 2 µm. **b, h,** Representative 3D reconstructions (Imaris *Spots* module) corresponding to **a** and **g**, showing Hsf1–Med15 (**b**) and Hsf1–Pol II (**h**) condensates in WT, *med5Δ,* and *med16Δ* cells after 10 min HS. Foci were detected using a fixed diameter of 0.48 µm. Light shading denotes Hsf1 puncta colocalized with Med15 (**b**) or Rpb3 (**h**). Scale bars, 0.5 µm. **c, i,** Percentage of cells containing ≥2 Hsf1–Mediator (**c)** or Hsf1–Pol II (**i**) co-condensates after 10 min HS. Data represent mean ± s.d. from two independent biological replicates. **d, j,** Violin plots showing the number of Hsf1, Mediator (**d**), and Hsf1, Rpb3 (**j**) puncta per cell after 10 min of heat shock, quantified using Imaris *Cells* module. Data from two independent biological replicates are shown. Horizontal lines indicate median values. **e, k,** Fraction of Hsf1 puncta colocalized with Med15 (**e**) or Pol II (**k**), quantified from 3D reconstructions using Imaris *Spots* module. Approximately 100 cells per strain were analyzed across two independent biological replicates. Data are mean ± s.d. **f, l,** Violin plots showing the distribution of the shortest volume-to-volume distances between Hsf1 and Med15 foci (f) or Hsf1 and Rpb3 foci (l) after 10 min HS in the indicated strains. Center lines indicate medians. Statistical significance was determined using a nonparametric two-tailed test for comparisons in **c, e, i, k** and one-way ANOVA for **d, f, j, l.** *, P < 0.05, **, P < 0.01, ***, P < 0.001.

In contrast, *MED16* deletion did not impair Hsf1 condensate formation, consistent with earlier observations (**Fig. 2**), but markedly reduced the fraction of cells containing Mediator and Pol II condensates (**Fig. 6a-c, g-i**), as well as the proportion of Hsf1 puncta colocalized with Mediator or Pol II (**Fig. 6e, f, k, l**). While the number of Hsf1 puncta per cell was unchanged in *med16Δ* cells, Mediator and Pol II puncta were substantially decreased (**Fig. 6d, j**), indicating that Med16 is required for efficient assembly of Mediator and Pol II condensates and for stable incorporation of transcriptional machinery into Hsf1 condensates. Notably, recruitment of Hsf1, Pol II, and Med15 to *HSR* genes was comparable between heat-shocked WT and *med16Δ* cells (**Fig. S15**), indicating that Med16 primarily regulates condensate organization rather than factor recruitment. Together, these findings support a hierarchical model in which Hsf1 condensate formation depends on the Mediator Tail triad, whereas productive transcriptional condensates, *HSR* gene coalescence, and robust transcriptional output additionally require Med16.

## Discussion

While Mediator has been shown to be a central component of transcriptional condensates in both lower and higher eukaryotes, how its individual subunits shape condensate organization, genome architecture, and transcription remains unclear. Here, we have investigated this question in *Saccharomyces cerevisiae* and identified a hierarchical organization of the Mediator Tail module that links IDR-driven condensate formation to higher-order chromatin architecture and robust transcription during the HSR.

### Mediator Tail triad subunits deploy their IDRs to drive Hsf1 condensate formation, 3D genome restructuring and *HSR* gene activation in acutely stressed cells

Our computational and genetic analyses identify Med2, Med3, and Med15 as an IDR-rich (**Fig. 1**), phase-separation–competent Tail triad that is essential for the formation of Hsf1 condensates (**Fig. 2**). These subunits exhibit extensive intrinsic disorder, enrichment of glutamine/asparagine residues, and high LLPS propensity—sequence features previously shown to promote condensate assembly through multivalent, weak interactions^37,38^. We found that loss of any Tail triad component markedly suppressed Hsf1 condensate formation and compromised *HSR* gene coalescence and transcription (**Figs. 2, 3).** Previous studies have shown that Med2-Med3-Med15 can physically interact with gene-specific transcription factors and be directly recruited to UAS regions^34,35,64–66^. Here, we have extended these findings by demonstrating that the IDRs within the Tail triad are functionally essential and promote Hsf1 condensate formation during the HSR. Notably, deletion or depletion of Med16, which disrupts the physical linkage between the Tail triad and core Mediator^35,66,70^, does not abolish Hsf1 condensate formation (**Fig. 2**). Instead, the released IDR-rich Tail triad is sufficient to interact with Hsf1 (suggested by Hsf1 ChIP analysis of *med16Δ* vs. WT; **Figs. S10** and **S15**), likely through multivalent IDR–IDR interactions, to enable heat shock–induced transcriptional activation of *HSR* genes (**Figs. 3d** and **S6).** These findings, together with the observation that deletion of either the ABD2 domain of Med15 or the C-terminal IDRs of Med2 / Med3 profoundly affects Hsf1 condensate formation, 3D genome architecture, and *HSR* gene activation (**Fig. 5)**, highlight the ability of the Tail triad to drive the three pillars of the heat shock transcriptional response, and do so via its IDRs. The graded phenotypes observed upon IDR deletion—less severe than complete subunit loss—suggest that these IDRs primarily tune condensate strength and transcriptional efficiency rather than serving as absolute on/off switches.

### Hsf1 condensate formation and genome restructuring can be mechanistically uncoupled

An important implication of this work is that transcriptional condensate formation and 3D genome organization are mechanistically separable. Deletion of any of the three IDR-rich Mediator tail triad subunits disrupts both Hsf1 condensate formation and *HSR* gene coalescence, consistent with a condensate-dependent mode of genome organization^16,31^. In contrast, loss of Med16 selectively abolishes intergenic and intragenic *HSR* gene interactions without impairing Hsf1 condensate assembly (**Figs. 2** and **3**). In this regard, the role of Med16 resembles that of two nuclear basket proteins, Mlp1 and Nup2, that are critical for driving HS-induced *HSR* gene coalescence yet dispensable for Hsf1 condensate formation^33^.

In addition, our data support a hierarchical regulatory model in which Hsf1–Mediator Tail triad interactions nucleate Hsf1 condensates through IDR-mediated assembly, while additional Mediator-dependent mechanisms orchestrated by the triad and Med16 stabilize Pol II– and Mediator-enriched transcriptional complexes and promote long-range chromatin contacts across *HSR* genes (model presented in **Fig. 7**). This functional separation is conceptually aligned with recent studies demonstrating that condensate formation and downstream transcriptional processes, including productive transcription and genome restructuring, can be genetically and mechanistically uncoupled^16,31,33,46^. Together, these findings establish Hsf1 condensate nucleation and genome organization as distinct yet coordinated layers of transcriptional regulation during the HSR.

**Figure 7.**
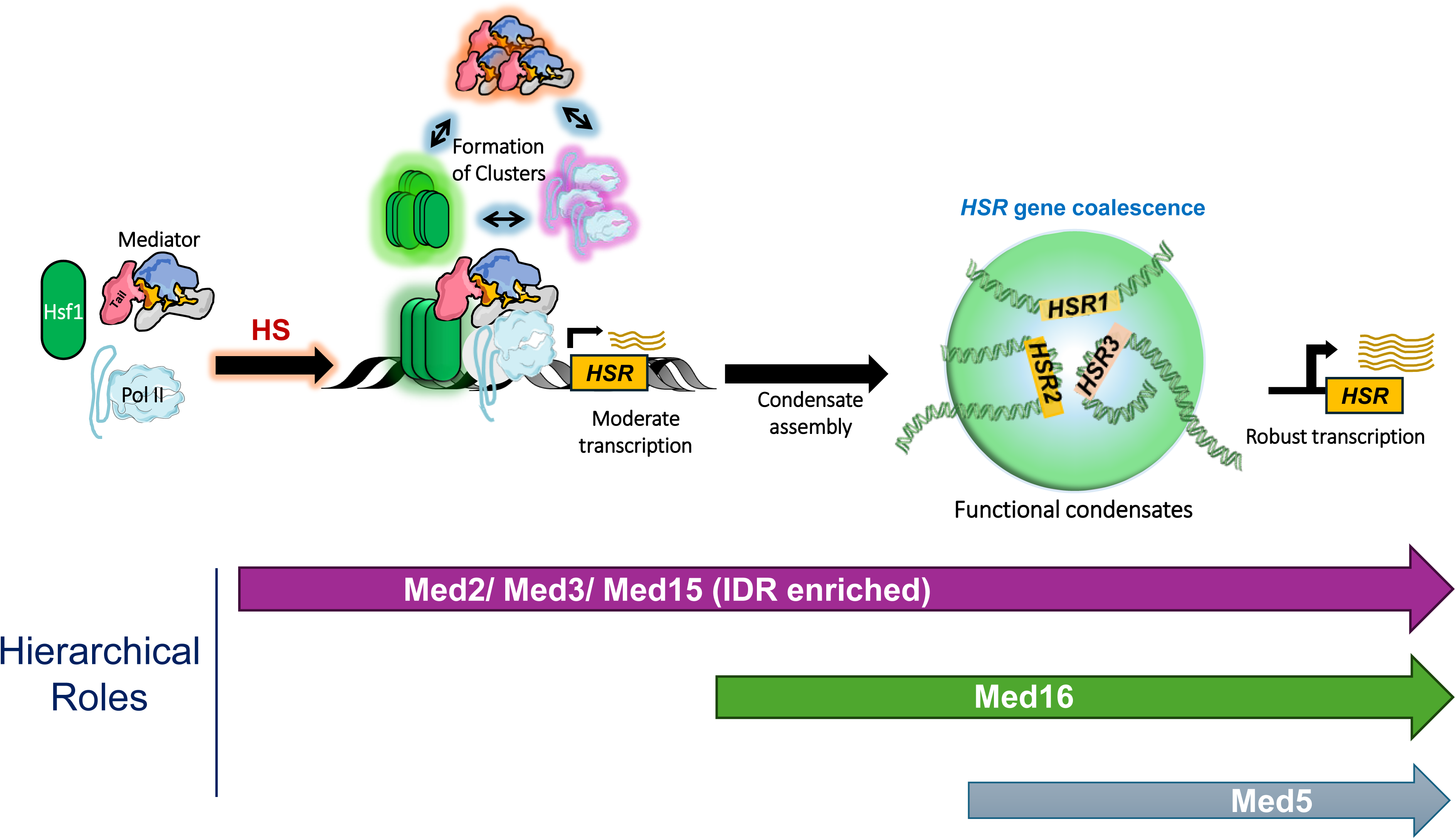
| Hierarchical model of Mediator Tail subunit regulation of the heat shock response. Upon acute heat shock (HS), Hsf1 trimerizes and, together with Mediator, Pol II and associated factors, is recruited to the upstream regulatory regions of *HSR* genes, activating their transcription. Hsf1, Mediator, and Pol II rapidly assemble into discrete intranuclear clusters and do so in a kinetically indistinguishable manner. Genetic analysis suggests that these subsequently mature into functional condensates that promote intergenic interactions among *HSR* gene loci, accompanied by enhanced levels of transcription. Within this framework, Mediator Tail subunits act hierarchically: the IDR–rich subunits Med2, Med3, and Med15 are essential for all steps; Med16 facilitates transcriptional condensate maturation and *HSR* gene activation and is essential for the global 3D restructuring of *HSR* genes; Med5 modestly enhances *HSR* interactions and *HSR* transcriptional output. Large multicolored arrows indicate the steps at which the indicated Tail subunits contribute.

### Med15 ABD2 integrates TF binding, condensate assembly, gene coalescence and transcription

Med15 has emerged as a central integrator of *HSR* regulation. Its activator-binding domains engage transcription factors through dynamic, “fuzzy” interfaces that have been shown to promote LLPS and transcriptional activation^57,58,60,61^. Our domain-specific analyses reveal that the intrinsically disordered ABD2 region plays a uniquely essential role, being required for Hsf1 condensate formation, *HSR* gene coalescence, recruitment of Mediator and Pol II, and robust transcriptional activation (**Figs. 4** and **S9**). Mechanistically, ABD2 enhances the occupancy of Hsf1, Mediator, and Pol II at *HSR* loci, thereby directly coupling condensate assembly to transcriptional output. These findings extend prior work demonstrating that Med15 mediates transcription through multivalent interactions with TFs^28,34,56,58,59,63^ by directly linking a specific Med15 IDR to condensate biology and genome organization. In contrast, other ABDs contribute more modestly, primarily influencing intrachromosomal architecture and factor occupancy, highlighting functional specialization among Med15 interaction surfaces. Whether the dominant role played by ADB2 is due to its being the primary target of Hsf1 was not addressed here. Other yeast and mammalian TFs have been observed to interact with the four ABDs in a “fuzzy” manner^57,58,60^ or primarily with the KIX domain^55^. Future work will address the question of which Med15 domain(s) Hsf1 physically contacts.

### Med16 promotes *HSR* gene coalescence by facilitating the formation of productive transcriptional condensates

Our live cell colocalization analyses uncover a distinct and essential role for Med16 in organizing the transcriptional machinery during the HSR. Med16 is specifically required for the formation and/or stabilization of Mediator and Pol II condensates and their incorporation into Hsf1 condensates (**Fig. 6**), while being dispensable for Hsf1 condensate assembly (**Fig. 2**). In the Med16 mutant, Hsf1 puncta form; however, these condensates fail to efficiently recruit or retain Mediator and Pol II, and this correlates with defective higher-order genome organization and reduced transcriptional output.

These observations indicate that Hsf1 condensate nucleation alone is insufficient to drive *HSR* gene coalescence and is consistent with the findings of others that implicate Pol II condensation as a key driver of *cis-* and *trans-HSR* intergenic interactions in HS cells^31^. Together, these results position Med16 as a critical organizer of transcriptional machinery, acting downstream of Hsf1 condensate formation to enable genome organization and robust transcription during stress.

### A hierarchical model for Mediator-driven transcriptional regulation

Together, our findings support a hierarchical and modular model of Mediator function during the HSR. In this framework, Hsf1 cooperates with the intrinsically disordered, phase-separation-prone Mediator Tail triad to nucleate transcriptional condensates through multivalent IDR-mediated interactions, consistent with current models of condensate-driven transcriptional activation^28,29,39,48,71^. Within this triad, Med15—particularly its intrinsically disordered ABD2 region—acts as a central integrator, linking transcription factor binding to condensate assembly and transcriptional activation. The C-terminal IDRs of Med2 and Med3 further reinforce condensate stability and enhance recruitment of Mediator and Pol II, thereby promoting robust transcriptional output.

In parallel, the structured Mediator subunit Med16 functions independently of Hsf1 condensate nucleation to organize Mediator–Pol II assemblies and facilitate higher-order chromatin interactions required for productive transcription. Med5 plays a detectable, albeit comparatively minor, role in promoting the formation of Hsf1 condensates, stimulating 3D genome restructuring, and contributing to HS-induced transcriptional activation (see **Figs. 7** and **S16** for a schematic model and graphic summary). This division of labor supports a model in which condensate nucleation, transcriptional machinery stabilization, and genome organization represent distinct yet coordinated regulatory layers. Given the evolutionary conservation of Mediator architecture^1,4,3,5^ and the widespread role of IDR-driven transcriptional regulation, these principles are likely to extend beyond the yeast HSR to diverse stress responses, enhancer-driven gene programs, and pathological transcriptional states in higher eukaryotes.

We note that a recent study observed no effect on heat shock-induced Hsf1 condensation when Med15 was conditionally depleted using the anchor-away (AA) technique^31^. A key difference between the AA approach and ours is that anchoring Med15 in the cytoplasm depletes the nucleus of not only Med15 but also potentially other factors (including other Mediator subunits^35^), in marked contrast to our subunit-specific approach, which does not (**Fig. S3g**). These contrasting observations imply that the Triad serves to counter an activity within one or more nuclear proteins that suppresses inappropriate Hsf1 condensation, an idea that will be tested in future work.

### Limitations and future directions

While our findings establish a central role for Mediator Tail subunits in coordinating transcriptional condensate formation, genome organization, and transcription during the HSR, several important questions remain. First, although computational analyses and genetic perturbations strongly implicate LLPS–driven mechanisms, direct biochemical reconstitution of the Mediator Tail and interactions of individual subunits with Hsf1 will be necessary to quantitatively define the properties of the condensates, constituent stoichiometries, and dynamic behavior. Second, our study focuses on an acute stress-responsive transcriptional program in yeast; whether the modular and hierarchical principles uncovered here extend to developmental enhancers, super-enhancers, or disease-associated regulatory elements in higher eukaryotes remains to be determined. Finally, although Med16 emerges as a key organizer of the transcriptional machinery and higher-order genome architecture, independent of Hsf1 condensate nucleation, the specific molecular interfaces and cofactors that mediate these functions remain unknown. One such candidate is Pol II, as cryo-EM analysis indicates that Med16 interacts with Rpb1 and Rpb9 in reconstituted Mediator-PIC complexes, an interaction that does not directly involve any other Tail subunit (S. Nagai and R.D. Kornberg, personal communication). Future studies integrating *in vitro* phase separation assays, high-resolution live-cell imaging—including single-particle tracking—and genome-wide chromatin conformation approaches such as Hi-C will be essential to elucidate how Mediator integrates condensate biology with 3D genome organization to regulate transcriptional output across diverse cellular contexts.

## Acknowledgements

We thank David Pincus and Surabhi Chowdhary for critical reading of the manuscript; Amoldeep S. Kainth for Med15 schematics; Steve Hahn, Tom Ellis and Surabhi Chowdhary for plasmid constructs; Kelly Tatchell for yeast knockout strains; and Shigeki Nagai and Roger D. Kornberg for sharing unpublished information. This work was supported by NIH R01GM13988 and an LSUHSC intramural grant awarded to DSG and Ike Muslow post– and pre-doctoral fellowships awarded to GM, SM and LSR.

## Methods

### Yeast Strain Construction

*Saccharomyces cerevisiae* Mediator subunit deletion mutants (*med2Δ, med3Δ, med5Δ, med15Δ* and *med16Δ*) and *MED15* domain deletion mutants were generated through homologous recombination using W303-1A and LRY037 as recipient strains^1^. PCR amplification of *med2Δ::KAN-MX* and *med15Δ::KAN-MX* was performed using chimeric primers of the respective genes and plasmid pFA-6a-KAN-MX6 as template. In contrast, *med3Δ, med5Δ* and *med16Δ* amplicons were amplified from genomic DNA isolated from the respective strains contained in the Res-Gen deletion collection. *MED15* domain deletion mutations were amplified from plasmid constructs^2^ (generous gift of Steve Hahn, Fred Hutchinson Cancer Center; **See Suppl. Table 2**) and subsequently transformed into either W303-1A or LRY037. Med15-13xMyc expressing strains were generated by homologous recombination with the Med15 C-terminal region PCR-amplified from ASK215 genomic DNA^3^. *MED3–3×FLAG* strains were generated by PCR amplification of the 3×FLAG cassette from pFA6a-3×FLAG using *MED3*-specific chimeric primers, followed by transformation into ASK218. *MED15* domain deletions were subsequently introduced into this background.

Degron-containing strains were generated by PCR amplification of the mini-AID*–9xMyc cassette from pHyg-AID*–9xMyc, pKan-AID*–9xMyc, or pNat-AID*–9xMyc (ref. ^4^) using chimeric primers homologous to the C-terminus of the target gene, followed by transformation into LRY016-derived strains^5^. For microscopy experiments, *RPB3-mCherry::hphMX6* and *MED15-mCherry::hphMX6* cassettes were amplified from SCY002 and SCY001, respectively^3^, and introduced into LRY037 to generate LRY039 and LRY040. Corresponding Mediator subunit deletion mutants (*med2Δ, med3Δ, med5Δ, med15Δ* and *med16Δ*) were subsequently constructed using LRY039 and LRY040 as parental strains. Yeast strains, plasmids and primers used in this study are listed in **Suppl. Tables 1–3**.

### CRISPR Editing

The intrinsically disordered regions (IDRs) of Med2 (residues 290-400) and Med3 (residues 301-374) were deleted using a CRISPR–Cas9 strategy following protocols developed by the Tom Ellis lab, Imperial College London^6^. Single-guide RNAs (sgRNAs) targeting the Med2 and Med3 IDRs were designed using the Benchling sgRNA design tool. Forward and reverse complementary sgRNA oligonucleotides were phosphorylated, annealed to generate double-stranded DNA, and cloned into the guide RNA assembly vector pWS082 using BsmBI-based Golden Gate assembly. Correct sgRNA insertion disrupts the GFP cassette on the vector backbone, enabling selection of non-fluorescent colonies. Purified sgRNA plasmids, verified by Nanopore sequencing, were digested with EcoRV and gel purified. The Cas9–sgRNA repair vector pWS173 was linearized by BsmBI digestion and gel-purified prior to yeast transformation.

Targeted deletion of the Med2 and Med3 IDRs was achieved by transforming strains W303-1A, LRY039 and LRY040 with a mixture of linearized Cas9–sgRNA repair vector, purified sgRNA-encoding DNA specific to Med2 or Med3, PCR-amplified donor DNA containing 100 bp homology arms upstream and downstream of the deleted IDR region, and salmon sperm DNA at a ratio of 100 ng: 200 ng: 3 μg: 50 μg, respectively. Pertinent sgRNA sequences, donor templates, primers, and plasmids are provided in **Suppl. Tables 2** and **3**.

### Yeast Culture and Heat Shock Treatment

For molecular biology experiments, cells were cultivated in rich YPDA medium (1% yeast extract, 2% peptone, 2% dextrose, and 20 mg/L adenine) at 30°C until reaching mid-logarithmic growth (OD_600_ = 0.6-0.8). Thermal stress (heat shock (HS)) was applied by rapidly elevating the culture temperature from 25° to 39°C through addition of an equal volume of 55°C YPDA medium, followed by incubation at 39°C for designated time points. The non-heat-shock (NHS) samples were mixed with an equal volume of 25°C YPDA and maintained at 25°C. Samples were kept at their respective temperatures using a water bath with constant shaking.

For experiments involving auxin-induced degradation, a stock of Indole-3-acetic acid (IAA, Sigma-Aldrich) was prepared in 100% ethanol to a final concentration of 100 mg/mL (570 mM). Cultures received IAA at a working concentration of 1 mM; incubation was performed at the time points indicated in the figures and text. To optimize degradation, samples were kept at 25°C for up to 1 hr. Heat shock experiments were performed by incubating cells for a predetermined length of time with IAA, followed by a thermal upshift to 39°C by adding 55°C media supplemented with 1 mM IAA to maintain a constant IAA concentration. The NHS sample was used as a control; for this, an equivalent volume of 25°C medium containing 1 mM IAA was combined with the culture.

### Growth Assays

For growth kinetics analysis, a single colony from either a wild-type or mutant strain was inoculated into 10 ml of sterile yeast peptone dextrose adenine (YPDA) medium and incubated for 16 hours at 30°C with orbital shaking to attain mid-exponential phase. Subsequently, cultures were diluted to OD_600_ = 0.2 and incubated at either 30°C or 39°C with constant orbital shaking (200 rpm) to assess temperature-dependent growth. OD_600_ measurements were taken at 1-h intervals for at least 8 h. Experiments were performed in biological duplicates, and mean OD_600_ values were calculated to construct growth curves illustrating the temporal relationship between optical density and time.

### Spot Dilution Assays

Spot dilution assays were performed as previously described^5^. In brief, overnight cultures were standardized to OD_600_ = 0.5, and 10-fold or 5-fold serial dilutions were prepared. Aliquots (∼4 μl) of each dilution were spotted onto YPDA plates using a 64-prong stainless steel replicator. Plates were incubated for 48 hours at 30°C or 72 hours at 24°C and 39°C to assess cell growth and viability.

### Chromatin Immunoprecipitation (ChIP)

ChIP assays were performed as described previously^7^. Briefly, 50 ml of mid-log-phase cultures (OD_600_ = 0.6-0.8) were treated with 1% formaldehyde for cross-linking after heat shock at 39°C for the indicated time points. Unreacted formaldehyde was quenched with 2.5 M glycine (final concentration, 375 mM). Cells were pelleted, lysed, sonicated, and clarified. Chromatin lysates (500 μL aliquots from a total volume of 2000 μL) were then immunoprecipitated with 1.5 μL anti-Hsf1 antiserum^8^, 1.5 μL of anti-Rpb1 antiserum^9^ or 2.5 μL anti-Myc mAb (9E10; Santa Cruz Biotechnology, Inc.) for 16 h at 4°C with gentle rotation. Antibody-bound chromatin was captured using Protein A Sepharose beads (GE Healthcare) by incubating 16 h at 4°C. DNA was subsequently purified using phenol-chloroform extraction. Locus-specific primers **(Suppl. Table 4)** were used to quantify ChIP DNA by quantitative PCR (qPCR) on a CFX384 Touch Real Time PCR System (Bio-Rad). Data was normalized to 20% input DNA, and the percentage of input was calculated accordingly.

### Fluorescence Microscopy and Image Analysis

Hsf1-mNeonGreen (Hsf1-mNG) puncta imaging was performed as previously described^1^. Cells were grown in synthetic complete dextrose (SDC) medium supplemented with 0.1 mg ml⁻¹ adenine at 30°C to early log phase, then adhered to a VaHEAT substrate (Interherence GmbH, Germany) coated with 100 µg ml⁻¹ Concanavalin A (ConA; Sigma-Aldrich) in ddH₂O. After removal of unbound cells and medium, fresh SDC medium was added, and the chamber was sealed with a coverslip to minimize evaporation. For mAID strains, mid-log phase cells were pretreated with either vehicle control or auxin for 1 h prior to attachment onto ConA-coated VaHEAT substrates, followed by imaging at 25° and 39°C. Imaging was performed using an Olympus Yokogawa CSU-W1 spinning disk confocal system equipped with Hamamatsu Fusion sCMOS cameras and controlled by cellSens Dimension software. Z-stacks of 11 planes (0.5 µm spacing) were acquired at the indicated time points using 488 nm excitation (10% laser power, 200 ms exposure) for mNeonGreen-tagged proteins and 561 nm excitation (15% laser power, 200 ms exposure) for mCherry-tagged proteins.

Hsf1–mNG puncta were quantified as described previously^1^. The percentage of cells containing Hsf1–mNG, Med15–mCherry, or Rpb3–mCherry foci (**Figs. 2a,b, 6a,g**) was determined using the “*Cells*” module in Imaris v10.2.0. This analysis included cells exhibiting ≥2 foci. During Imaris segmentation, the nuclear diameter and spot diameter were set to 2 µm and 0.48 µm, respectively^10^. For Hsf1/Med15 and Hsf1/Rpb3 colocalization analysis (**Fig. 6 e, f, k, l**), foci were detected using the Imaris “*Spots*” module with a spot diameter of 0.48 µm. Colocalization was assessed by measuring the shortest volume-to-volume distance between spots, and foci were considered colocalized when this distance was ≤0.45 µm. Image cropping and 3D reconstruction were performed using FIJI/ImageJ (v1.53t)^11^. All quantified data were plotted using GraphPad Prism v10.

### Taq I Chromosome Conformation Capture (Taq I – 3C)

Taq I – 3C was performed as previously described^5,12^. Briefly, samples were crosslinked with 1% formaldehyde (15 min), quenched with 0.363 M glycine (10 min), and lysed for 40 min using glass beads in FA Lysis Buffer. Lysates were aliquoted and subjected to Taq I digestion (20 U/μL, 7 h, 60°C) and Quick T4 DNA ligation (10,000 cohesive units, 2 h, 25°C). Samples were then treated with RNase A (30 ng/μL, 40 min, 37°C) and Proteinase K (70 ng/μL, 16 h, 65°C), followed by phenol-chloroform extraction, ethanol precipitation, and qPCR quantification using locus-specific primers (**Suppl. Table 5)**.

### Reverse Transcription-qPCR (RT-qPCR)

RT-qPCR was performed as described^5^ using 25 mL cell culture aliquots. Cells were metabolically arrested and lysed using glass beads and acidic phenol-chloroform, followed by RNA extraction using acidic phenol-chloroform. Residual DNA was removed (Turbo DNA-free kit), and cDNA was synthesized (High-Capacity cDNA Reverse Transcription Kit, both from Applied Biosystems). cDNA was quantified on a CFX384 System using gene-specific primers (**Suppl. Table 6**) and a standard curve generated from genomic DNA. Quantified DNA was normalized to *SCR1*.

### Immunoblot Analysis

Immunoblot analysis was performed as previously described^5^. Briefly, log-phase cells were metabolically arrested with 20 mM sodium azide for 30 seconds, lysed, and proteins precipitated with trichloroacetic acid. Protein pellets were dried out and resuspended in Thorner buffer (8 M urea, 5% SDS, 40 mM Tris-HCl pH 6.8, 0.1 mM EDTA, 0.4 mg/mL Bromophenol Blue, 1% 2-Mercaptoethanol). Samples were neutralized with 2 M Tris base, incubated at 42°C for 15 minutes, and separated on SDS 4-20% polyacrylamide TGX precast gels (Bio-Rad). Proteins were transferred to PVDF membranes (Amersham, 0.2 μm) (or nitrocellulose, Bio-Rad, for AID experiments), blocked with 5% nonfat milk in TBST (10 mM Tris-HCl, pH 8, 150 mM NaCl, 0.05% Tween20) overnight at 4°C, and probed with primary antibodies (Myc, 1:1000, Santa Cruz sc-40; H3, 1:1000, Abcam ab1719; Pgk1, 1:10,000, ThermoFisher 459250) and secondary HRP-conjugated antibodies (1:5000, Santa Cruz) for 1 h. Protein bands were detected using Enhanced Chemiluminescence (ECL) reagent (Thermo Scientific 34580).

### Intrinsic Disorder, Structural Confidence and Phase Separation Analyses

Protein sequences were retrieved in FASTA format and analyzed using complementary sequence-based and machine-learning approaches to quantify intrinsic disorder and liquid–liquid phase separation (LLPS) propensity. Intrinsic disorder was predicted using IUPred3 (long mode)^13^ and PONDR (VSL2)^14^ to generate per-residue disorder probabilities; residues with scores ≥0.5 were classified as disordered. The fraction of disordered residues per protein was calculated to determine the intrinsic disorder region (IDR) abundance. LLPS propensity was evaluated using FuzDrop (probability of spontaneous LLPS, pLLPS)^15^ and PhasePred^16^, extracting PS-self and PS-part scores for each protein. All the graphs were generated using GraphPad Prism (v10).

Structural confidence profiles were obtained from AlphaFold3^17^ models, and per-residue predicted Local Distance Difference Test (pLDDT) scores were used as an orthogonal measure of structural order, with regions of pLDDT <70 considered indicative of structural flexibility/disorder.

Intra-molecular interaction maps were generated using the FINCHES online^18^ tool (Mpipi-GG method; window size = 31). Amino acid composition was calculated using ExPASy ProtParam^19^. Default parameters were used for all analyses unless otherwise specified.

### Statistical Analysis

Statistical analyses were performed using GraphPad Prism, as specified in the figure legends. Differences between individual means were assessed using two-tailed, unpaired *t* tests assuming equal variance. For violin plot analyses, statistical significance was determined by one-way ANOVA.

