## Supplemental Figures for "Mediator Tail Subunits Hierarchically Couple Transcriptional Condensates to Gene Activation and Genome Organization"

### Supplementary Figure S1

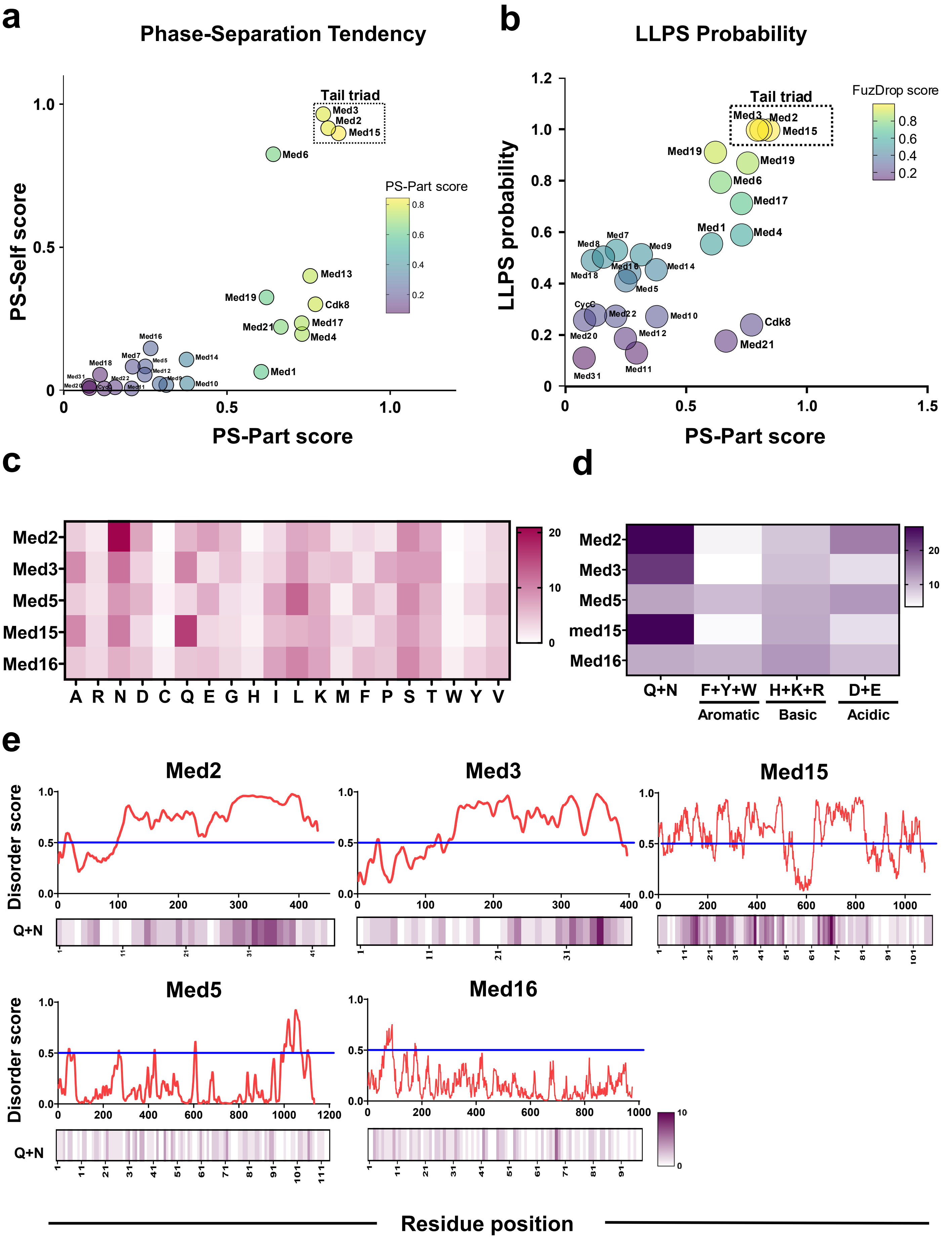

**Figure S1 | The Mediator Tail triad has enhanced phase-separation tendencies.**

**a,** Phase separation propensity of Mediator subunits predicted by PhaSePred, shown as PS-part (partner-dependent phase separation) and PS-self (self-phase separation) scores; the Tail triad (Med2, Med3 and Med15) is highlighted.

**b,** Comparison of phase separation propensity predicted by PhaSePred (PS-part score) and liquid–liquid phase separation probability predicted by FuzDrop for the indicated Mediator subunits.

**c,** Heatmap illustrating the percentage amino acid composition of each Tail subunit.

**d,** Comparative heatmap depicting the relative abundance of selected amino acid classes in Mediator Tail subunits, including glutamine and asparagine (Q + N), aromatic residues (Phe + Trp + Tyr), basic residues (His + Lys + Arg), and acidic residues (Asp + Glu).

**e,** Composite maps showing location of predicted intrinsically disordered regions based on IUPred3 analysis (top) aligned to distribution of glutamine (Q) and asparagine (N) residues (bottom) across the five Tail subunits. Q and N enrichment was calculated using a 10–amino acid sliding window and visualized as heatmaps along the protein sequence.

### Supplementary Figure S2

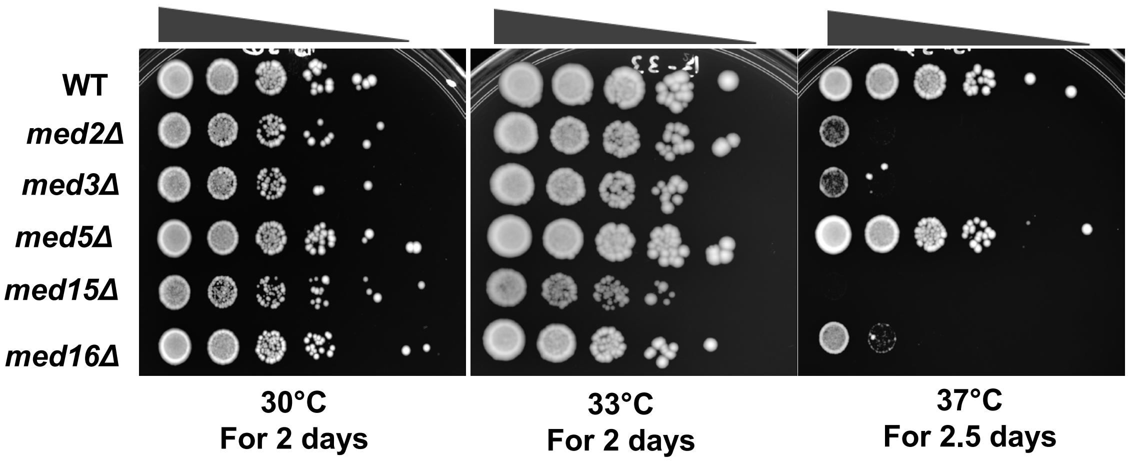


**Figure S2 | All Mediator Tail subunits except Med5 are required for growth at elevated temperature.** Tenfold serial dilutions of the indicated yeast strains were spotted onto YPDA medium and incubated at the indicated temperatures for 2.0 – 2.5 days, then photographed.

### Supplementary Figure S3

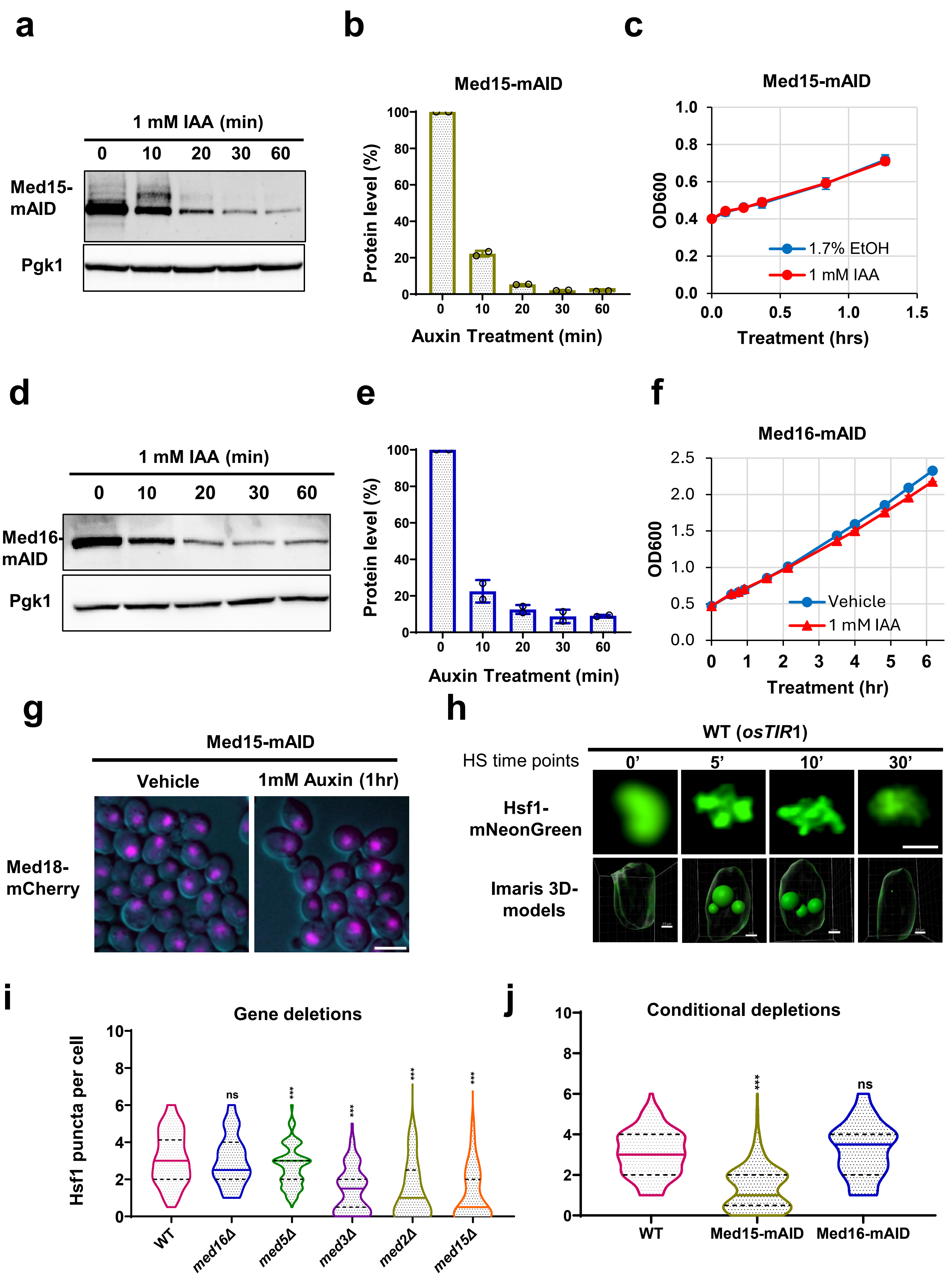

**Figure S3 | Characterization of Med15-mAID and Med16-mAID strains.**

**a, d,** Immunoblot analysis of Med15-mAID-9Myc and Med16-mAID-9Myc expression levels in cells treated with 1 mM indoleacetic acid (IAA) for the indicated times. Blots were probed with anti-Myc antibody (9E10) and Pgk1 as a loading control.

**b, e,** Quantification of Med15-mAID-9Myc and Med16-mAID-9Myc expression levels relative to Pgk1 at each time point, represented as the average of two independent experiments ± standard deviation.

**c, f,** Growth kinetics of the Med15-mAID and Med16-mAID strains in YPDA liquid culture. Cells were grown at 30°C, and optical density ($A_{600}$) was monitored over time.

**g,** Representative fluorescence micrographs of Med15-mAID cells expressing Med18–mCherry treated with vehicle or 1 mM IAA for 1 h at 25°C. Scale bar, 5 µm.

**h,** Representative fluorescence micrographs and corresponding three-dimensional reconstructions (Imaris v10.2.0, Cells module) of Hsf1 condensates in wild-type (LRY109) cells during a 39 °C heat-shock time course. Scale bar, 2 µm. Scale bar for Imaris-rendered 3D models, 0.5 µm.

**i, j,** Violin plots showing the distribution of Hsf1 puncta per cell after a 10 min HS, quantified using Imaris. Solid horizontal lines indicate median values; discontinuous lines depict quartiles. Statistical significance relative to WT was assessed by one-way ANOVA (*, $P < 0.05$; **, $P < 0.01$; ***, $P < 0.001$).

### Supplementary Figure S4

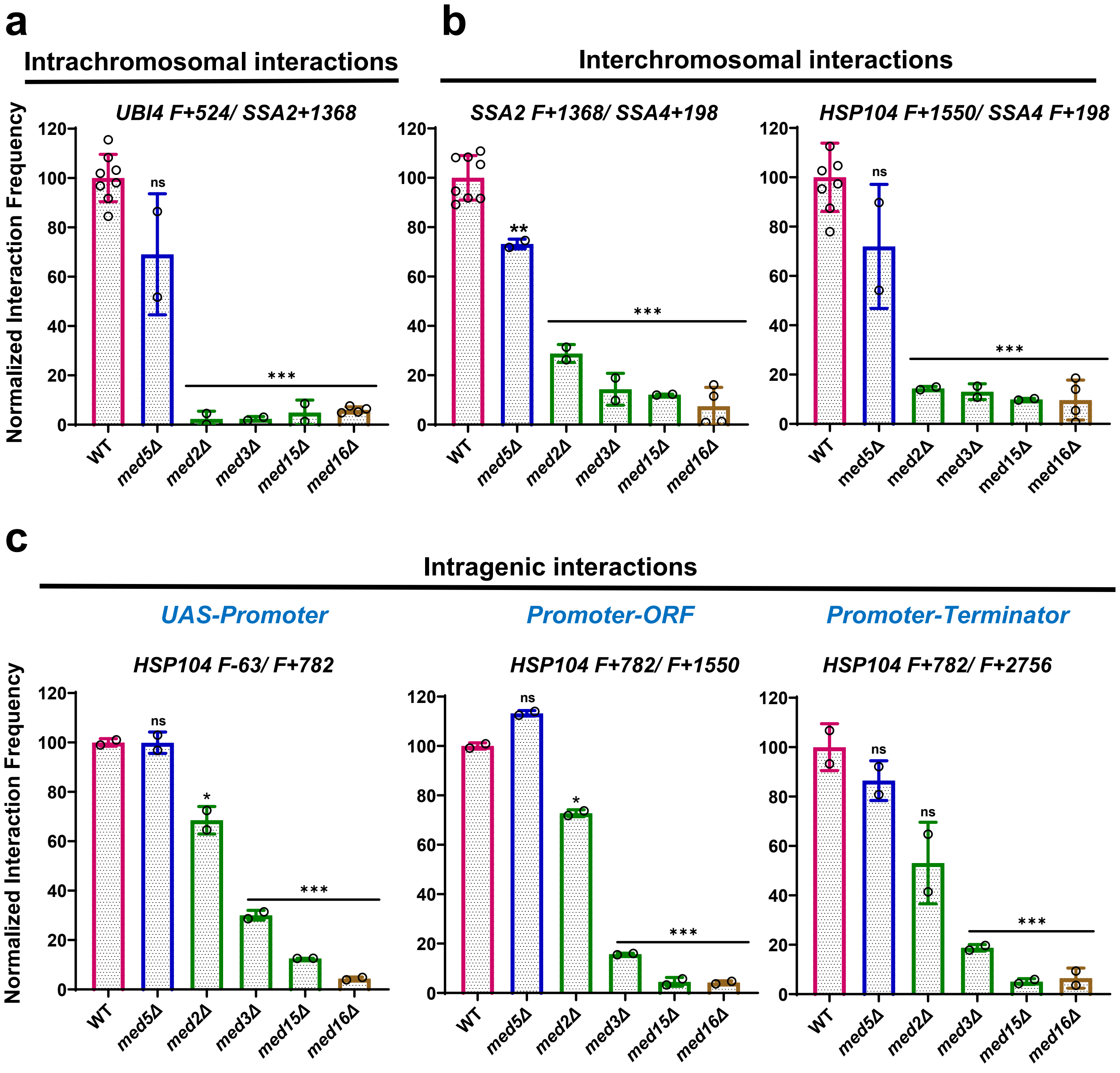

**Figure S4 | Mediator Tail subunits are required for heat-shock-induced *HSR* gene interactions.**

Shown are normalized interaction frequencies of *Heat Shock Response* (*HSR*) genes measured by Taq I - 3C in the indicated strains after 5 min of heat shock. **a** shows intrachromosomal (*cis*), **b** interchromosomal (*trans*), and **c** intragenic interactions. Green bars denote strains lacking a Tail Triad subunit. Data are presented as mean ± s.d. from the number of independent samples indicated. Statistical significance relative to wild-type (WT) was assessed using an unpaired two-tailed *t*-test (*, $P < 0.05$; **$P < 0.01$; ***$P < 0.001$).

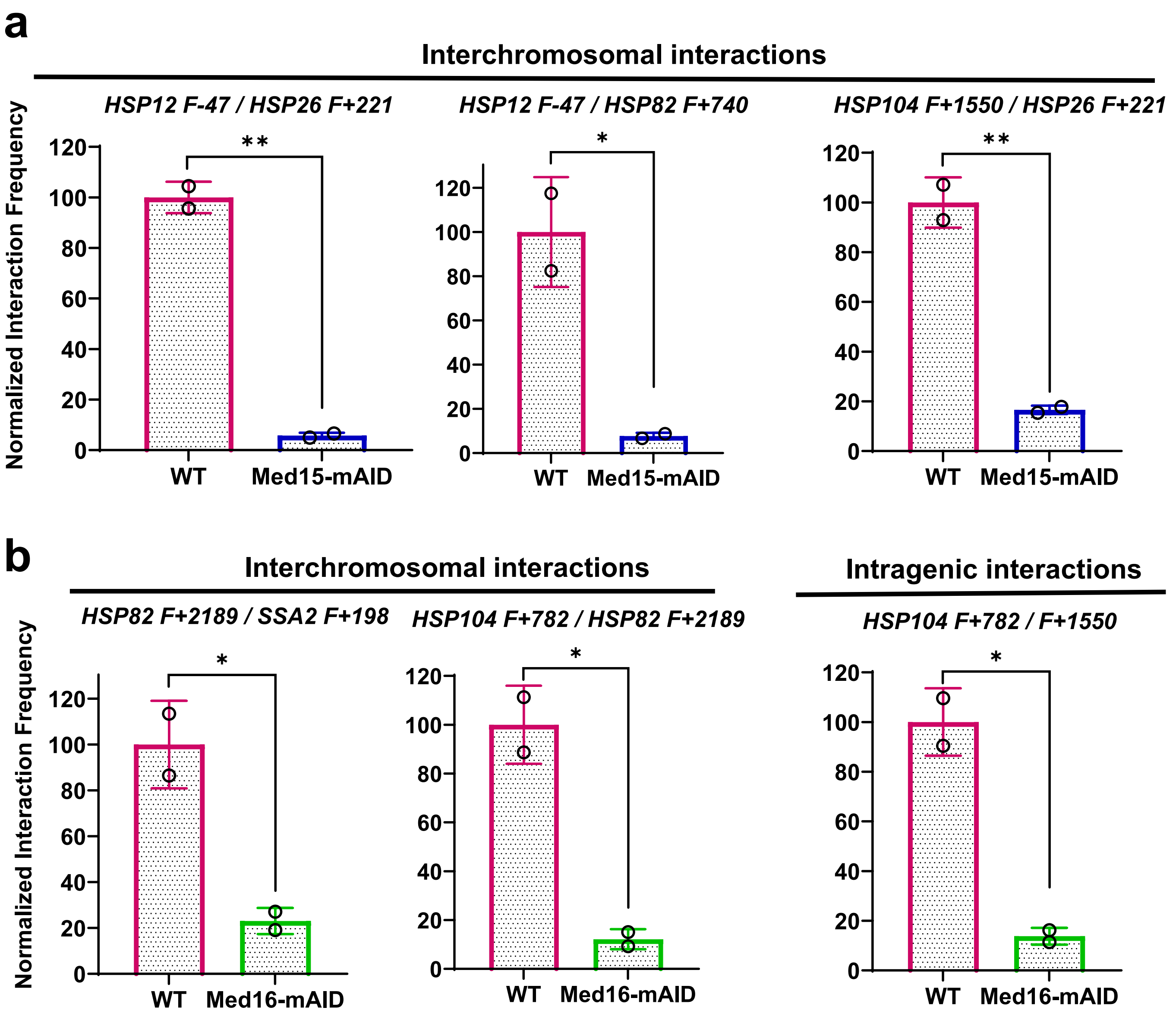


**Figure S5 | Conditional depletion of either Med15 or Med16 disrupts heat-shock-induced interactions between *HSR* genes.**

Effects of conditional depletion of Med15 (**a**) and Med16 (**b**) on *Heat Shock Response* (*HSR*) gene interactions assessed by Taq I - 3C. Cells were pretreated with 1 mM IAA for 60 min at 30 °C prior to heat shock at 39°C, and 3C analysis was performed after 2.5 min HS. Data represent mean ± s.d. of two independent experiments. Statistical significance relative to WT was determined using unpaired two-tailed *t*-tests. *, $p < 0.05$; **, $p < 0.01$; ***, $p < 0.001$.

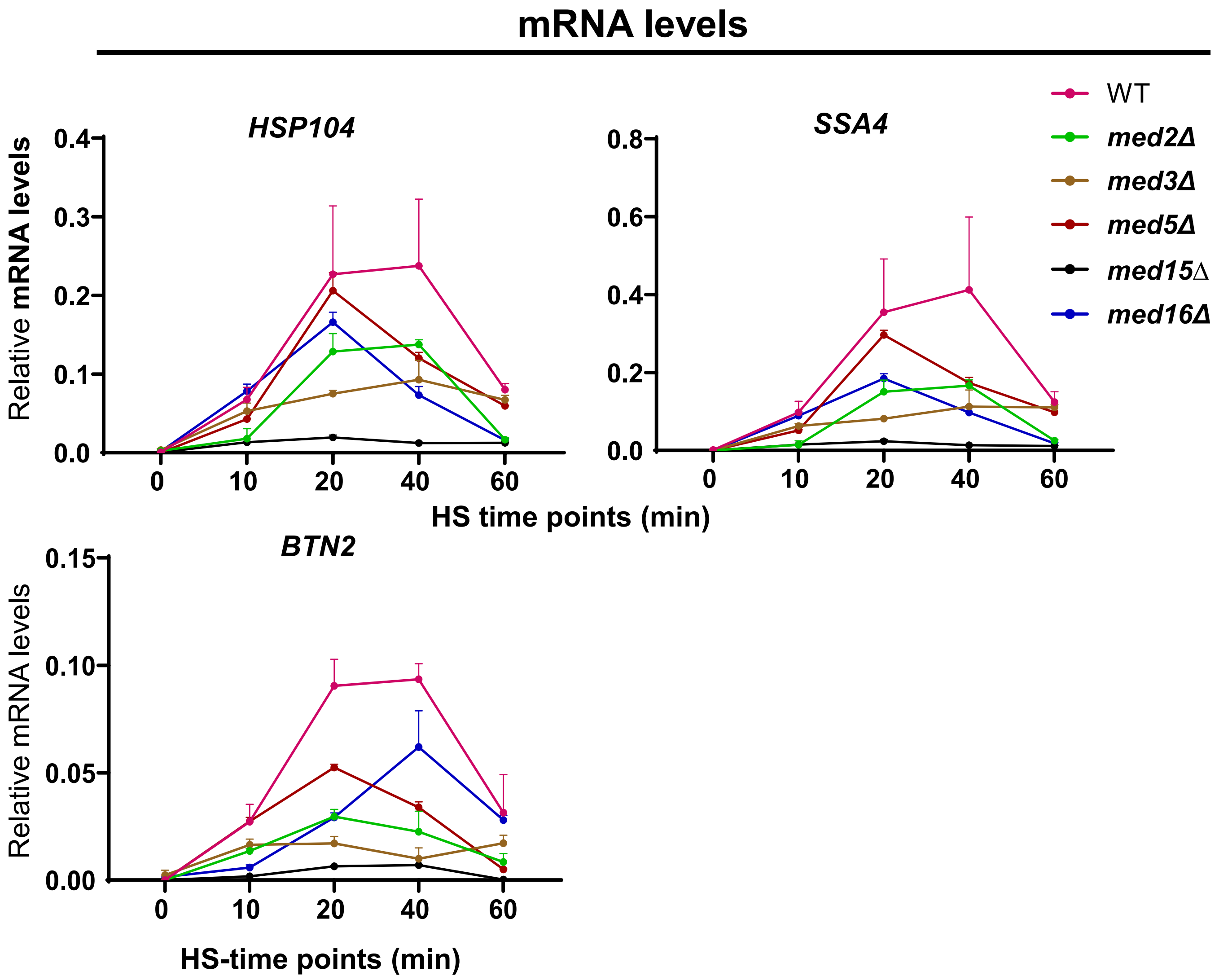


**Figure S6 | Loss of any Tail triad subunit – Med2, Med3 or Med15 – substantially impairs *HSR* mRNA accumulation in HS-induced cells.** RNA levels of *HSP104*, *SSA4* and *BTN2* were determined by RT–qPCR. Data represent mean ± s.d. from two independent biological replicates for each strain, except for WT, for which *n* = 6. Statistical significance relative to WT was assessed using an unpaired two-tailed *t*-test. *, $p < 0.05$; **, $p < 0.01$; ***, $p < 0.001$.

#### Supplementary Figure S7

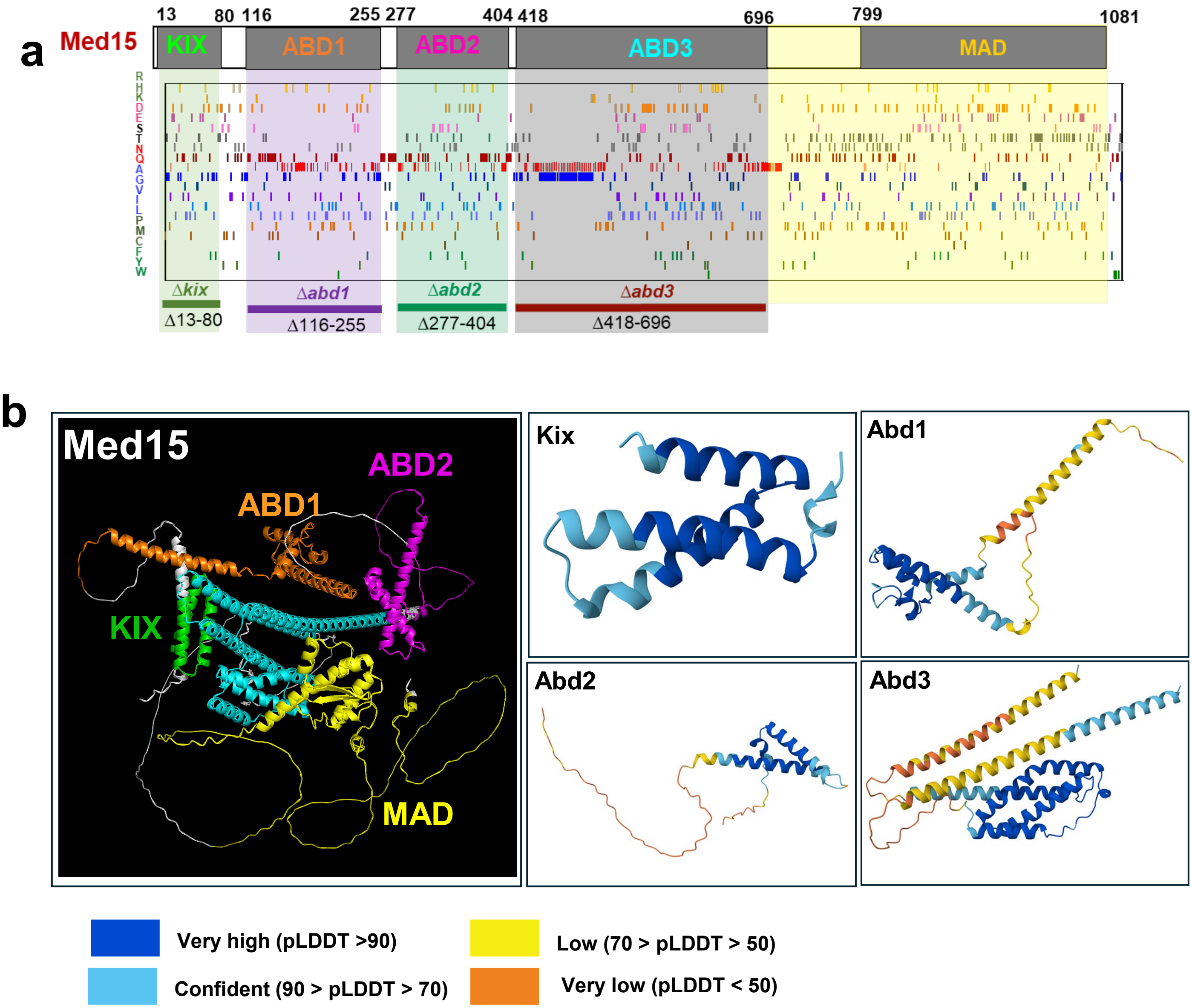


**Figure S7 | Med15 contains both structured and intrinsically disordered regions.**

**a,** Domain architecture of Med15 showing the positions of four activator-binding domains (ABDs) required for Med15 function. MAD, Mediator Associated Domain. Bottom, amino acid composition of individual Med15 domains.

**b,** AlphaFold3-predicted structures of Med15. The left panel shows the predicted structure of full-length Med15, and the right panels show the predicted structures of individual ABDs. Colors indicate prediction confidence, as shown in the scale below.

#### Supplementary Figure S8

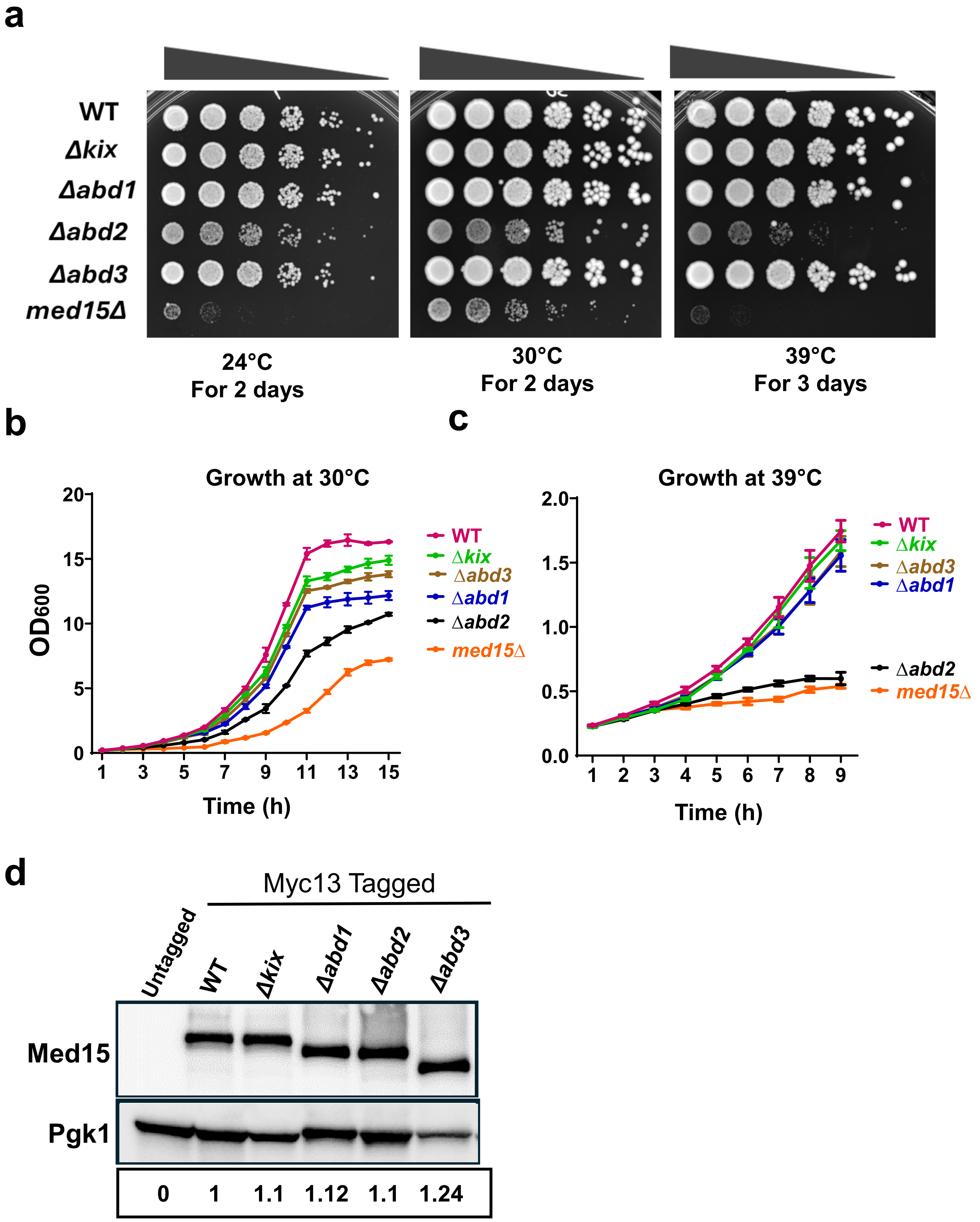

**Figure S8 | The Med15 ABD2 domain is required for growth fitness at elevated temperature.**

**a,** Tenfold serial dilution analysis of the indicated strains spotted on YPDA medium and incubated at the indicated temperatures for the specified times.

**b, c,** Growth kinetics of WT and Med15 domain-deletion strains cultured at 30 °C (**b**) and 39 °C (**c**). Data represent mean ± s.d. from two independent biological replicates.

**d,** Immunoblot analysis of Myc13-tagged Med15 domain-deletion constructs probed with anti-Myc antibody. Pgk1 was used as a loading control. Relative protein expression levels are indicated.

#### Supplementary Figure S9

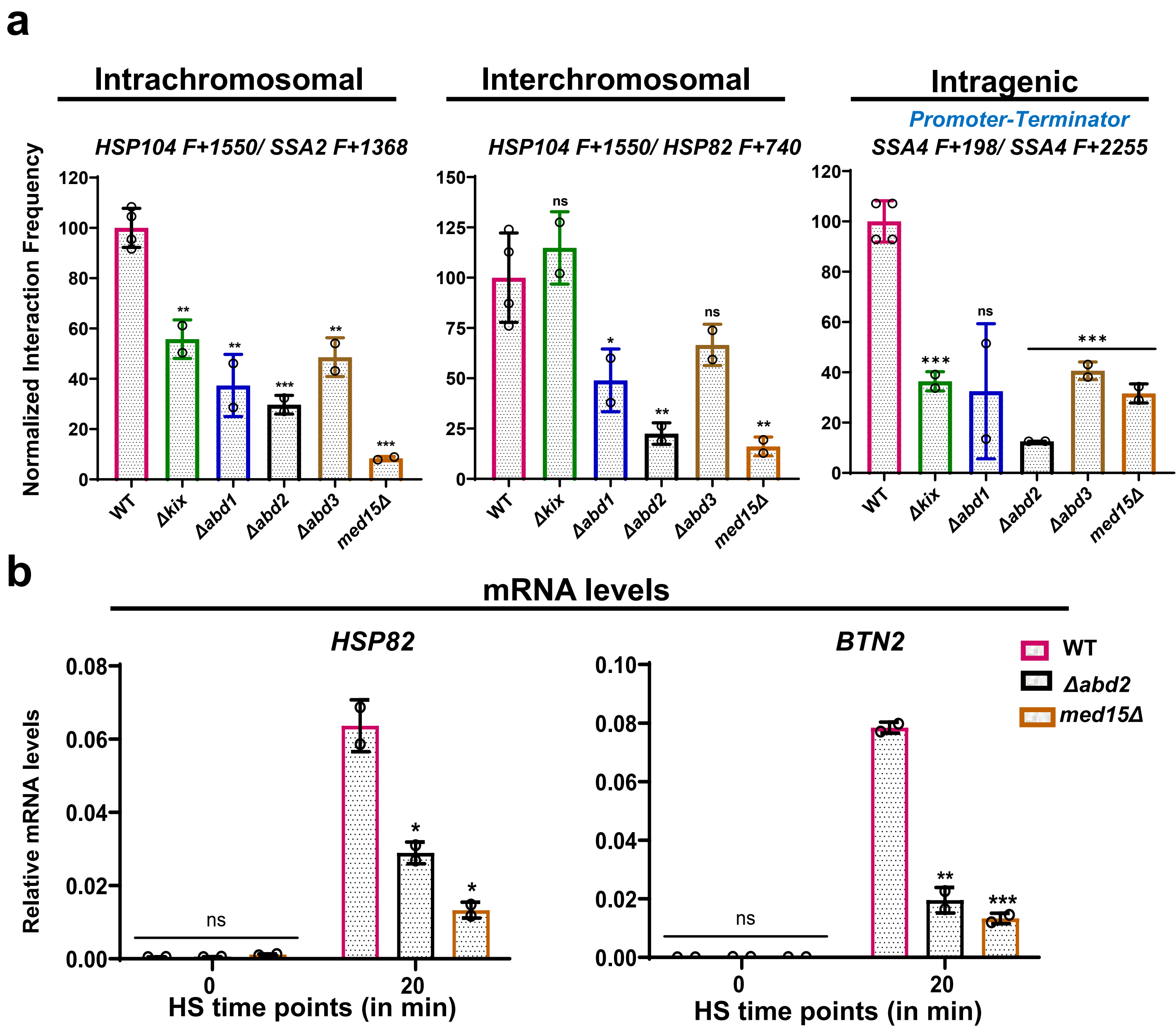


**Figure S9 | The Med15 ABD2 domain plays an important role both in driving 3C interactions and in promoting *HSR* gene expression in acutely stressed cells.**

**a,** Normalized interaction frequencies of *Heat Shock Responsive* genes measured by Taq I - 3C in the indicated strains after 5 min of heat shock. Data represent mean ± s.d. from the indicated number of independent samples. Statistical significance relative to WT was assessed using an unpaired two-tailed *t*-test. *, $p < 0.05$; **, $p < 0.01$; ***, $p < 0.001$.

**b,** Relative mRNA levels of representative *HSR* genes determined by RT–qPCR. Data represent mean ± s.d. from two independent biological replicates. Statistical significance was determined as above.

### Supplementary Figure S10

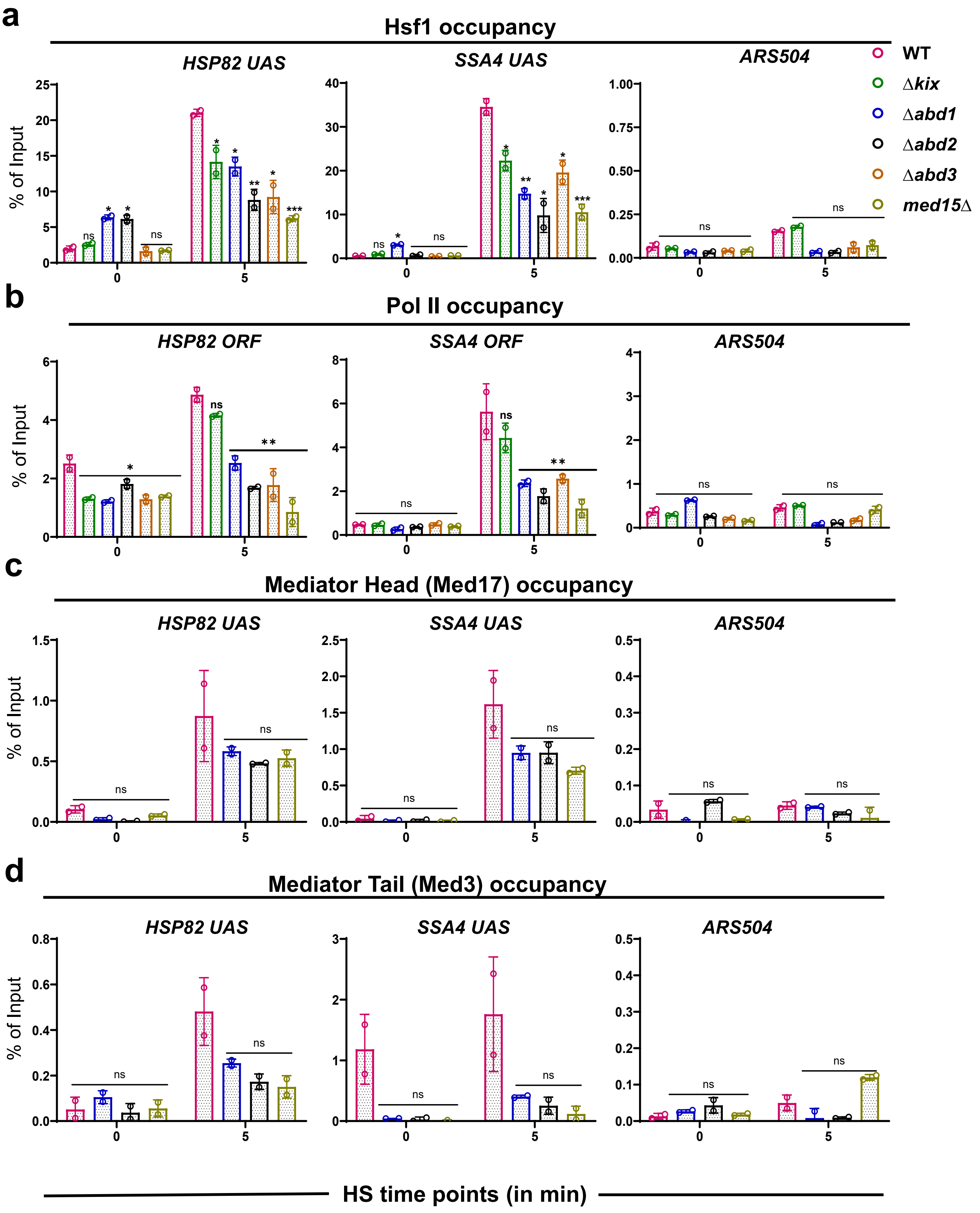

**Figure S10 | Med15 activator-binding domains promote recruitment of Hsf1, Pol II and Mediator to *Heat Shock Response* genes, with ABD2 playing the central role.**

Chromatin immunoprecipitation analysis of **a** Hsf1, **b** RNA polymerase II (Rpb1), **c** Mediator Head subunit Med17–Myc9, and **d** Mediator Tail subunit Med3–Flag occupancy at the *HSP82*, *SSA4*, and *ARS504* (control) loci in WT, *med15Δ* and Med15 domain-deletion strains. Cells were maintained at 24°C (0 min) or subjected to heat shock at 39°C for 5 min. Data represent mean ± s.d. from two independent biological replicates ($n = 2$). Statistical significance relative to WT was assessed using an unpaired two-tailed *t*-test. *, $p < 0.05$; **, $p < 0.01$; ***, $p < 0.001$; ns, not significant.

### Supplementary Figure S11

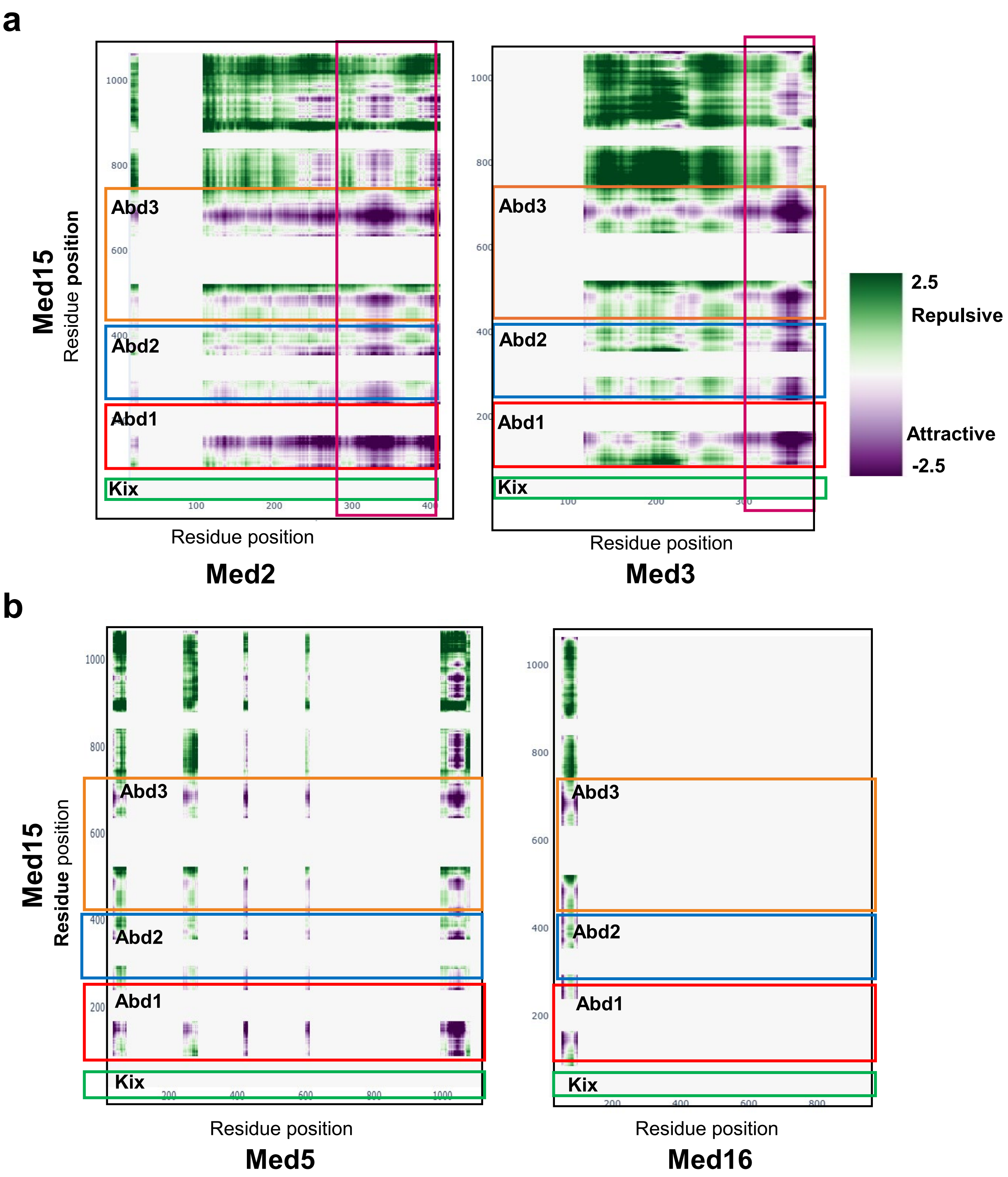

**Figure S11 | Med2 and Med3 C-terminal IDRs are predicted to interact strongly, while Med5 and Med16 are predicted to interact weakly, with Med15 activator-binding domains (ABDs).**

**a,** FINCHES-predicted inter-residue interaction maps between Med15 activator-binding domains (KIX and ABD1–ABD3; highlighted, vertical axis) and the C-terminal intrinsically disordered regions (IDRs) of Med2 and Med3 (highlighted; horizontal axis). Attractive interactions are shown in purple, and repulsive interactions are shown in green.

**b,** FINCHES-predicted inter-residue interaction maps between Med15 ABDs and the indicated regions of Med5 and Med16.

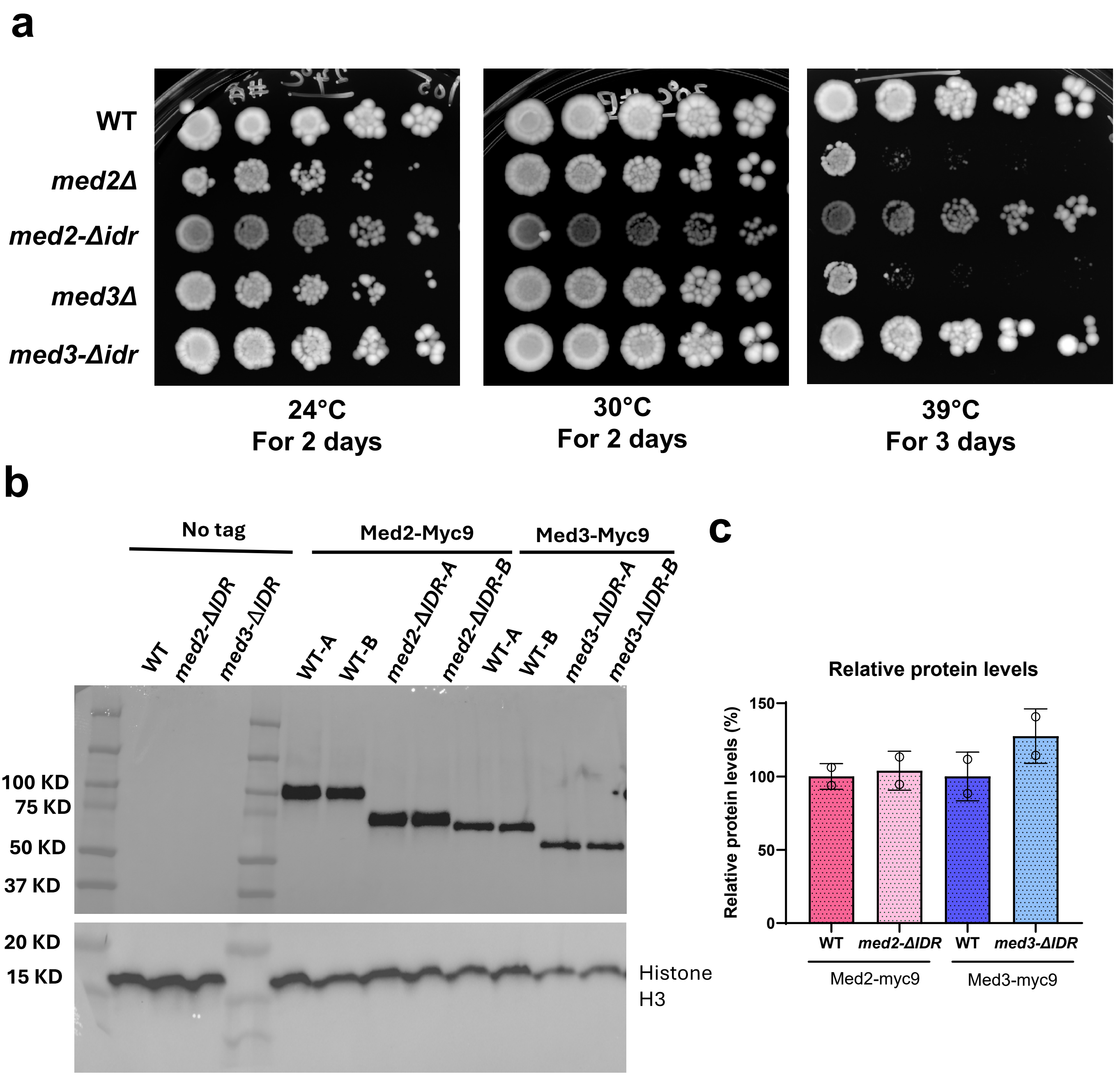


**Figure S12 | Med2 and Med3 IDRs contribute to growth fitness at elevated temperature.**

**a,** Fivefold serial dilution analysis of the indicated strains spotted on YPDA medium and incubated at the indicated temperatures to assess growth fitness and viability under heat-stress conditions.

**b,** Immunoblot analysis of the indicated Myc9-tagged strains probed with anti-Myc antibody. Histone H3 was used as a loading control.

**c,** Quantification of relative protein expression levels shown in **b**, presented as bar graphs. Data represent results from two independent biological replicates.

### Supplementary Figure S13

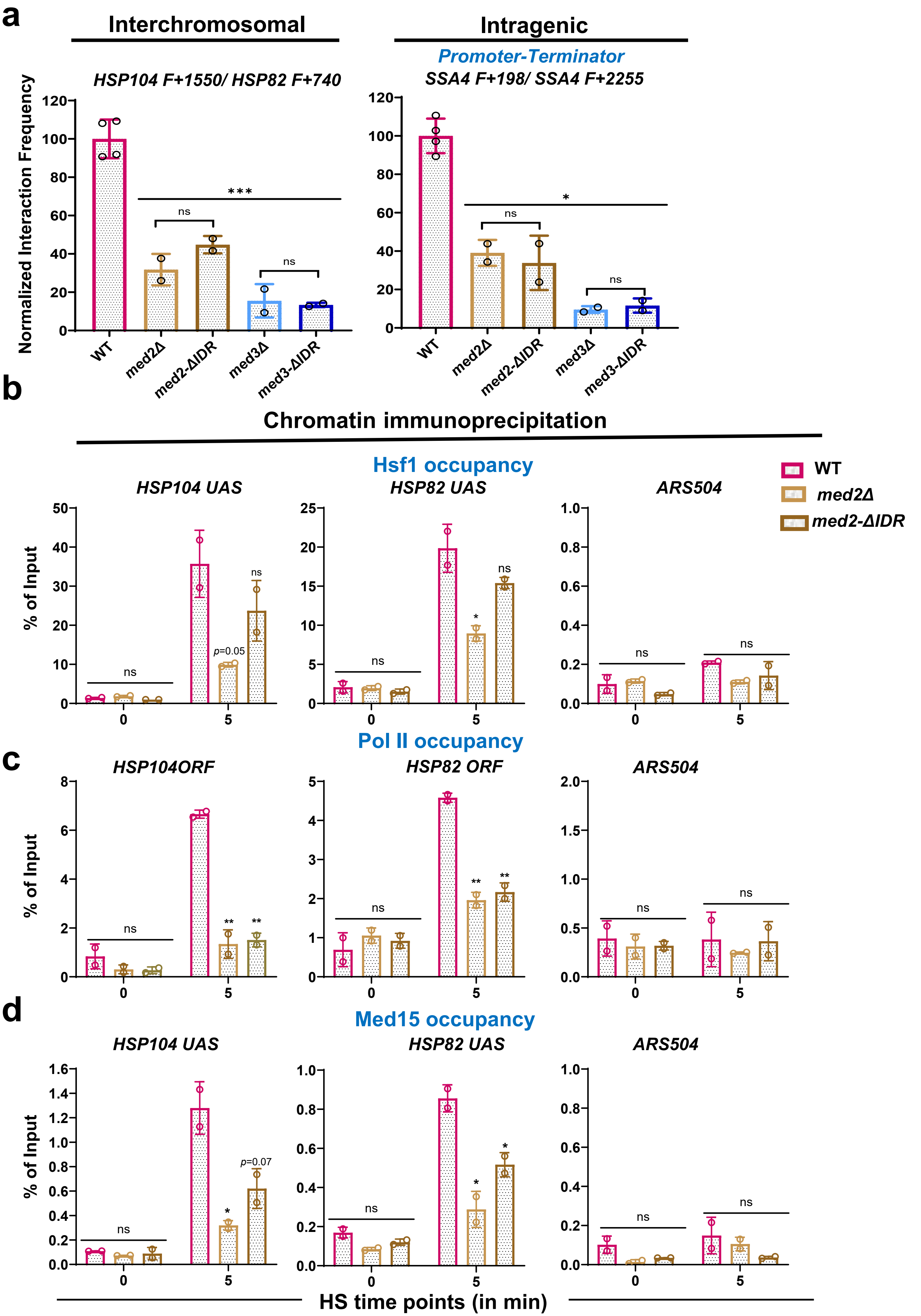

**Figure S13 | Med2 and Med3 C-terminal IDRs regulate the heat-shock response by modulating genome topology and Hsf1, Mediator and Pol II occupancy.**

**a,** Normalized interaction frequencies of *Heat Shock Responsive* genes measured by Taq I - 3C after 5 min of heat shock. Data represent mean ± s.d. Statistical significance relative to WT was assessed using an unpaired two-tailed *t*-test (*, $p < 0.05$; **, $p < 0.01$; ***, $p < 0.001$ ).

**b–d,** ChIP analysis of **b** Hsf1, **c** RNA polymerase II (Rpb1), and **d** Med15–Myc13 occupancy at the *HSP104*, *HSP82*, and *ARS504* (control) loci under basal conditions (24 °C) or after heat shock at 39 °C for 5 min. Data represent mean ± s.d. of two independent biological replicates. Statistical significance was determined as above.

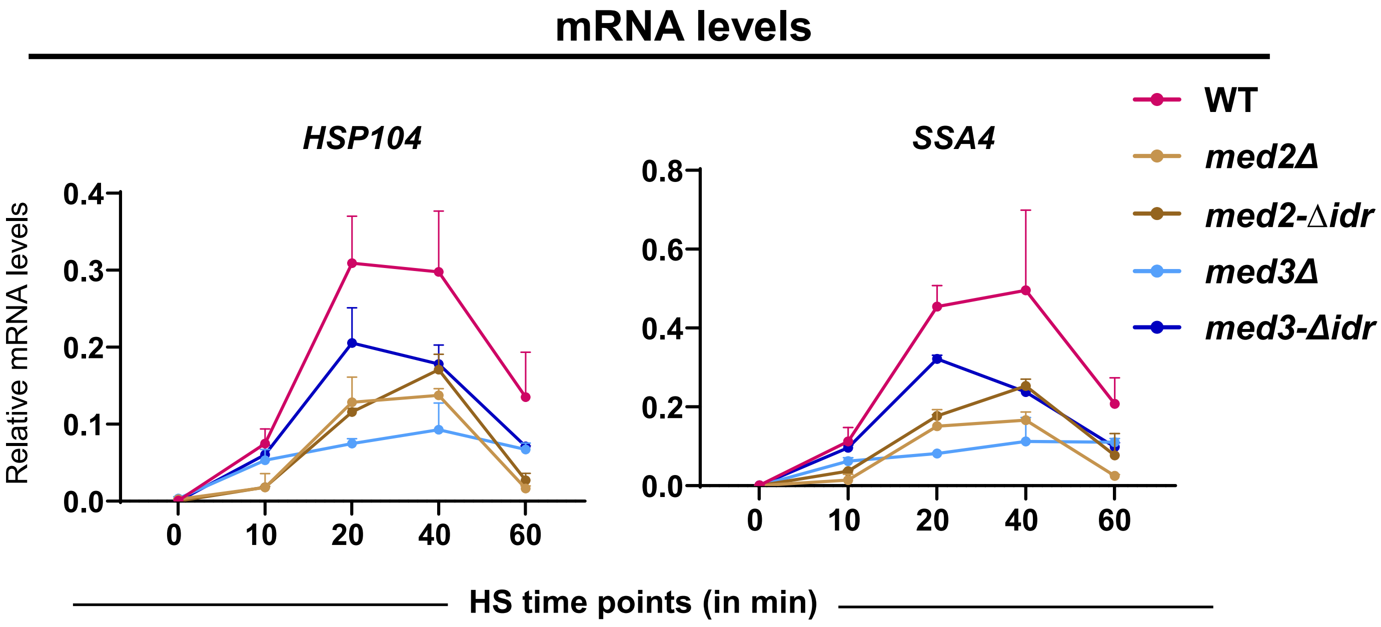


**Figure S14 | Loss of intrinsically disordered regions in Med2 and Med3 impair *HSR* gene expression.**

Relative mRNA levels of *HSP104* and *SSA4* determined by RT–qPCR. Data represent mean ± s.d. from two independent biological replicates for each strain, except for WT, for which $n = 4$. Statistical significance relative to WT was assessed using an unpaired two-tailed *t*-tests: *, $p < 0.05$; **, $p < 0.01$; ***, $p < 0.001$.

### Supplementary Figure S15

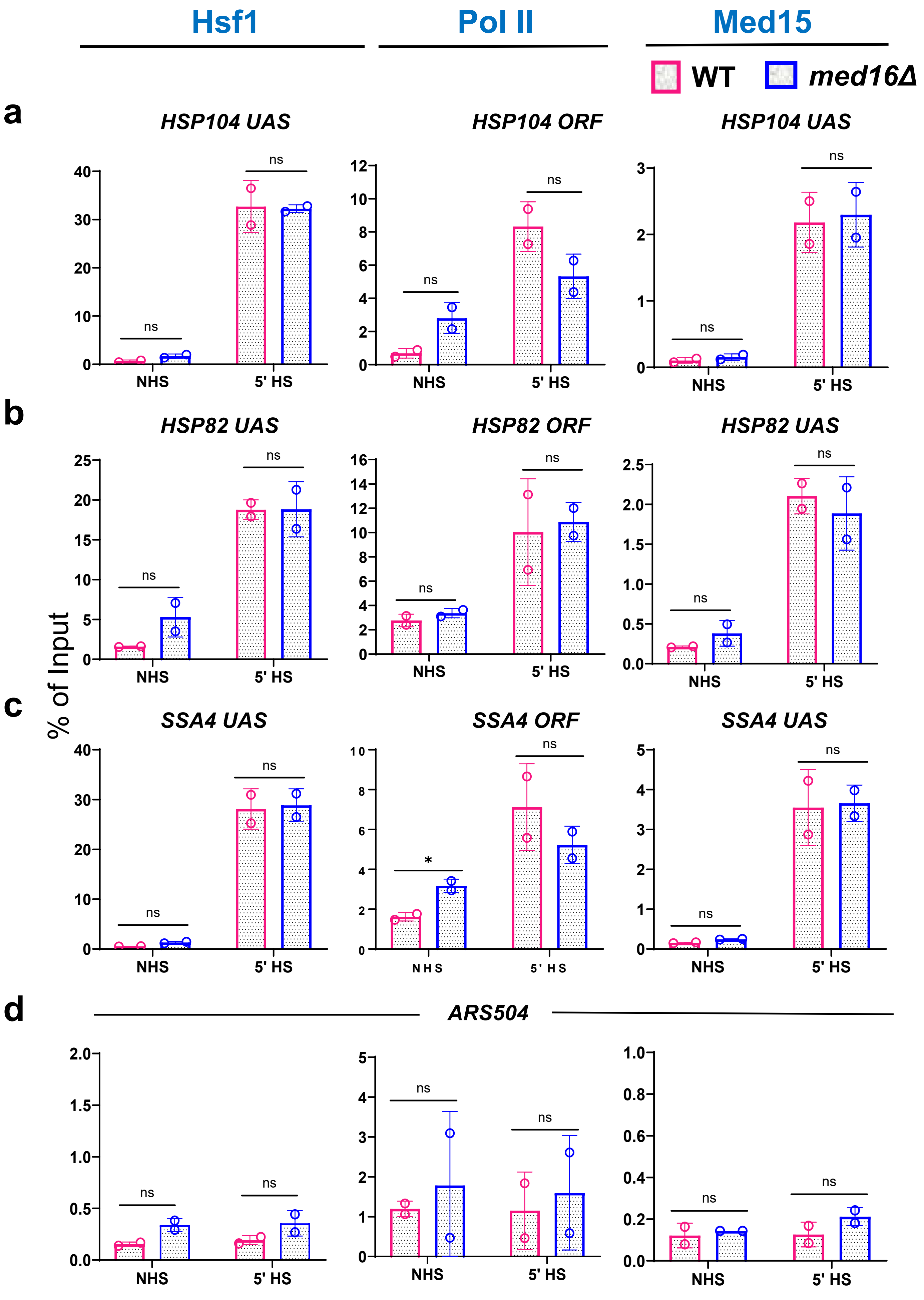

**Figure S15 | Deletion of Med16 has no detectable effect on the recruitment of Hsf1, Pol II, or Mediator to *HSR* genes.**

**a–d,** ChIP analysis of **b** Hsf1, **c** RNA polymerase II (Rpb1), and **d** Med15–Myc13 occupancy at the *HSP104*, *HSP82*, and *ARS504* (control) loci under basal conditions (24 °C) or following heat shock at 39 °C for 5 min. Data represent mean ± s.d. from two independent biological replicates. Statistical significance was assessed using an unpaired two-tailed *t*-test. *, $p < 0.05$; **, $p < 0.01$; ***, $p < 0.001$ (WT vs. *med16*Δ).

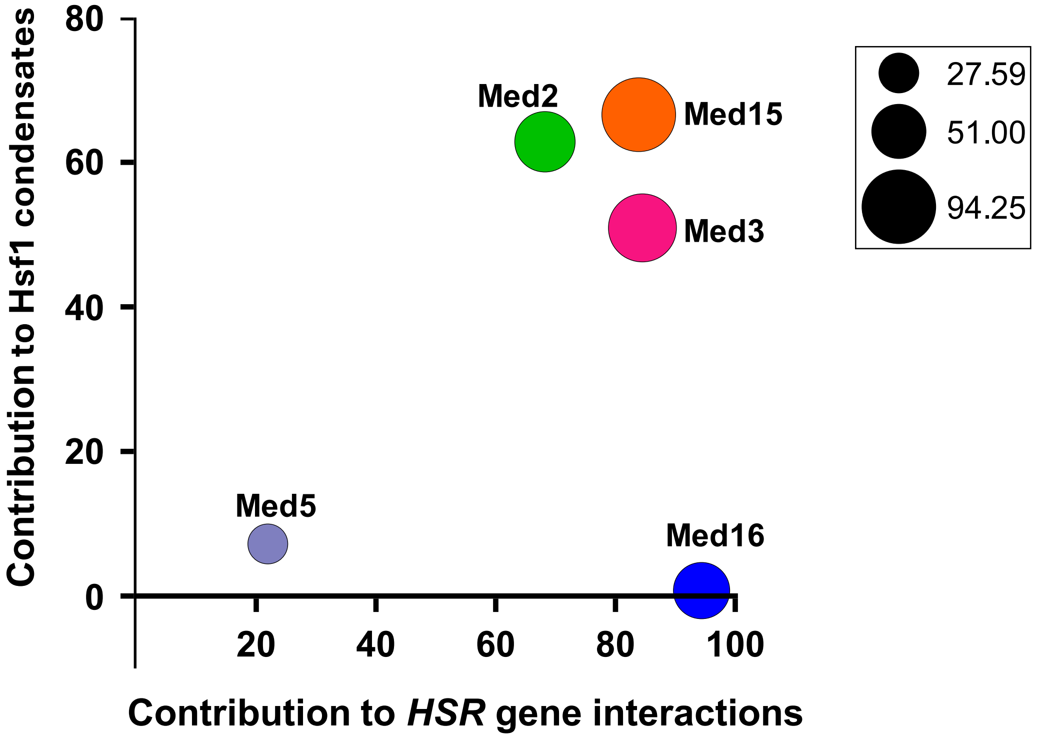


**Figure S16 | Hierarchical contributions of Mediator Tail subunits to the heat shock response.**

Bubble plot summarizing the relative roles of Mediator Tail subunits in regulating the HSR through control of Hsf1 condensate formation, *HSR* gene interactions, and *HSR* gene transcription. The x-axis shows the loss of normalized interaction frequency (relative to the isogenic WT strain) between *HSP104 (Chr. XII)* and *SSA4 (Chr. V*) following a 5 min HS due to loss of the indicated subunit. The y-axis similarly indicates the relative contribution of each subunit to Hsf1 condensate formation following a 10 min HS. Bubble size represents the contribution of each subunit (% loss relative to WT level) in *HSP104* mRNA abundance following a 20 min heat shock.
