## Supplemental Tables for "Mediator Tail Subunits Hierarchically Couple Transcriptional Condensates to Gene Activation and Genome Organization"

**Table 1. Yeast Strains**

| Strain Name | Genotype | Source \| Reference |
|---|---|---|
| W303-1A | *MATa ade2-1 trp1-1 can1-100 leu2-3,112 his3-11,15 ura3-1* | R. Rothstein |
| W303-1B | *MATα ade2-1 trp1-1 can1-100 leu2-3,112 his3-11,15 ura3-1* | R. Rothstein |
| GMY001 | LRY037; *med15Δ::KAN-MX* | This study |
| GMY005 | W303-1A; *med15Δ::KAN-MX* | This study |
| GMY013 | LRY037; *med15-Δkix::URA3* | This study |
| GMY014 | W303-1A; *med15-Δkix::URA3* | This study |
| GMY024 | LRY037; *med15-Δabd1::URA3* | This study |
| GMY028 | W303-1A; *med15-Δabd1::URA3* | This study |
| GMY030 | LRY037; *med15-Δabd2::URA3* | This study |
| GMY034 | W303-1A; *med15-Δabd2::URA3* | This study |
| GMY036 | LRY037; *med15-Δabd3::URA3* | This study |
| GMY038 | W303-1A; *med15-Δabd3::URA3* | This study |
| GMY056 | ASK218; *MED3-3xFLAG::HIS3* | This study |
| GMY058 | GMY056; *med15-Δabd1::URA3* | This study |
| GMY059 | GMY056; *med15-Δabd2::URA3* | This study |
| GMY066 | GMY056; *med15Δ::KAN-MX* | This study |
| GMY067 | W303-1A; *MED15-13xMyc::HIS3* | This study |
| GMY068 | GMY014; *MED15-13xMyc::HIS3* | This study |
| GMY069 | GMY028; *MED15-13xMyc::HIS3* | This study |
| GMY070 | GMY034; *MED15-13xMyc::HIS3* | This study |
| GMY071 | GMY038; *MED15-13xMyc::HIS3* | This study |
| GMY076 | LRY037; *med2Δ::KAN-MX* | This study |
| GMY077 | LRY037; *med3Δ::KAN-MX* | This study |
| GMY078 | LRY037; *med5Δ::KAN-MX* | This study |
| GMY079 | LRY037; *med16Δ::KAN-MX* | This study |
| GMY081 | W303-1A; *med2Δ::KAN-MX* | This study |
| GMY082 | W303-1A; *med3Δ::KAN-MX* | This study |
| GMY083 | W303-1A; *med5Δ::KAN-MX* | This study |
| GMY084 | W303-1A; *med16Δ::KAN-MX* | This study |
| ASK215 | *MED15-13xMyc::HIS3* | Chowdhary et al, 2022 [1] |
| ASK218 | W303-1A; *MED17-9xMyc::TRP1* | This study |
| ASK220 | W303-1A; *med15-Δkix::URA3 MED17-9xMyc::TRP1* | This study |

| | | |
|---|---|---|
| SMY172 | LRY016; *HSF1-mNeonGreen::HIS3* | Mohajan et al, 2025 [2] |
| SMY256 | W303-1A; *med16Δ::KAN-MX* | This study |
| SCY001 | *MATa ADE2 trp1-1 can1-100 leu2-3,112 his3-11,15 ura3-1 HSF1-mVenus::HIS3 MED15-mCherry::hphMX6* | Chowdhary et al, 2022 [1] |
| SCY004 | *MATa ADE2 trp1-1 can1-100 leu2-3,112 his3-11,15 ura3-1 HSF1-mVenus::HIS3 RPB3-mCherry::hphMX6* | Chowdhary et al, 2022 [1] |
| LRY016 | W303-1B*; LEU2::pGPD1-osTIR1* | Rubio et al, 2024 [3] |
| LRY037 | W303-1B; *HSF1-mNeonGreen::HIS5* | Rubio et al, 2024 [3] |
| LRY039 | LRY037; *MED15-mCherry::hphMX6* | This Study |
| LRY040 | LRY037; *RPB3-mCherry::hphMX6* | This Study |
| LRY101 | LRY016, *MED15-mAID-9xMyc:: hphMX6* | This study |
| LRY103 | LRY016, *MED14-mAID-9xMyc::KAN-MX* | This Study |
| LRY104 | LRY016, *MED16-mAID-9xMyc::KAN-MX* | This study |
| LRY107 | LRY103; *RPB3-mCherry::hphMX6* | This study |
| LRY108 | LRY104; *RPB3-mCherry::hphMX6* | This study |
| LRY109 | SMY172; *RPB3-mCherry::hphMX6* | This study |
| LRY111 | LRY107; *HSF1-mNeonGreen::HIS3* | This study |
| LRY112 | LRY108; *HSF1-mNeonGreen::HIS3* | This study |
| LRY114 | LRY109; *MED15-mAID::NAT-MX* | This study |
| GMY086 | LRY039; *med3Δ:: KAN-MX* | This study |
| GMY087 | LRY039; *med5Δ:: KAN-MX* | This study |
| GMY088 | LRY039; *med16Δ:: KAN-MX* | This Study |
| GMY089 | LRY040; *med2Δ::KAN-MX* | This study |
| GMY090 | LRY040*; med5Δ:: KAN-MX* | This study |
| GMY091 | *LRY040; med16Δ:: KAN-MX* | This Study |
| GMY092 | W303-1A; *med2-ΔIDR* (Δ290-400) | This study |
| GMY093 | LRY039; *med2-ΔIDR* (Δ290-400) | This study |
| GMY094 | LRY040; *med2-ΔIDR* (Δ290-400) | This Study |
| GMY095 | W303-1A; *med3-ΔIDR* (Δ301-374) | This study |
| GMY096 | LRY039; *med3-ΔIDR* (Δ301-374) | This study |
| GMY097 | LRY040; *med3-ΔIDR* (Δ301-374) | This Study |
| GMY102 | W303-1A*; MED2-9xMyc::TRP1* | This study |
| GMY103 | GMY092*; med2-ΔIDR (Δ290-400)- 9xMyc::TRP1* | This study |
| GMY104 | W303-1A; *MED3-9xMyc::TRP1* | This Study |
| GMY105 | GMY095; *med3-ΔIDR*(Δ301-374) *-9xMyc::TRP1* | This study |
| GMY106 | GMY081; *MED15-13xMyc::HIS3* | This study |
| GMY107 | GMY092*; MED15-13xMyc::HIS3* | This Study |
| GMY110 | GMY084; *MED15-13xMyc::HIS3* | This Study |
| GMY111 | SMY256; *MED15-13xMyc::HIS3* | This Study |

**Table 2. Plasmids**

| Plasmid Name | Feature | Source \| Reference |
|---|---|---|
| pHyg-AID*-9myc | miniAID-9xMYC-HYGR | Morawska & Ulrich, 2013 [4] |
| pKan-AID*-9myc | miniAID-9xMYC-KANR | Morawska & Ulrich, 2013 [4] |
| pNat-AID*-9myc | miniAID-9xMYC-NATR | Morawska & Ulrich, 2013 [4] |
| pFA6a-KAN-MX6 | pFA6a-KAN-MX6 | Addgene Plasmid #39296 (Bahler et al, 1998) [5] |
| pWZV87 | 9xMYC-KITRP1 | Cosma et al, 1999 [6] |
| pFA6a-3xFLAG | 6xGLY-3xFLAG-HIS3-MX6 | Addgene plasmid #20753; Funakoshi & Hochstrasser, 2009 [7] |
| pEH30-Δ1 | pRS316; *GAL11* Δ13-80 | Herbig et al, 2010 [8] |
| pEH30-Δ4 | pRS316; *GAL11* Δ116-255 | Herbig et al, 2010 [8] |
| pEH30-Δ5 | pRS316; *GAL11* Δ277-404 | Herbig et al, 2010 [8] |
| pEH30-Δ6 | pRS316; *GAL11* Δ418-696 | Herbig et al, 2010 [8] |
| pWS082 | sgRNA entry vector | Tom Ellis (Addgene plasmid #90516); Shaw et al, 2019 [9] |
| pWS158 | Cas9 + sgRNA scaffold | Tom Ellis (Addgene plasmid #90517); Shaw et al, 2019 [9] |

**Table 3. Primers Used for Strain Construction**

| Name | Sequence (5' → 3') | Purpose |
|---|---|---|
| *MED2* Chimeric FP | GTTTAAATTTGAAGGCGGATCCTCCCAAATAAACTGCCCGTCTGAAAGTAATTTAGGTGACACTATAG | Amplification of *med2Δ::KAN-MX* cassette |
| *MED2* Chimeric RP | GAATGCACAACACGGTTTACAAGTCAATAGTTAACAATAGGAAGACCAAGTAATACGACTCACTATAGGG | |
| *MED2* conf FP | GAGGGTTGCTGGAATATTCACC | Confirmation of *med2Δ::KAN-MX* insertion |
| *MED2* conf RP | GAGATGCAGAGAGTTGAACAAGC | |
| *MED3* FP | GTCACCAAGACTTCTTCAATTAGG | Amplification of *med3Δ::KAN-MX* |
| *MED3* RP | GTTGTTCTGACATGGTCTGTCG | |
| *MED3* conf FP | GAAGTCGAACGACGAAGAAGC | Confirmation of *med3Δ::KAN-MX* insertion |
| *MED3* conf RP | GATGATATCTACACCACCATCCTGG | |
| *MED5* amp FP | CTATTCGTGGCTAGTATATGACC | Amplification of *med5Δ::KAN-MX* |
| *MED5* amp RP | GGAGTTAATGTCTATCTTTAATCTGC | |
| *MED5* conf FP | CAGGTCGAAGACTCCATCC | Confirmation of *med5::KAN-MX* insertion |
| *MED5* conf RP | GAAGTCACTCGTGAAGGTAGC | |
| *MED16* amp FP | CAGACCTGACCTTCTGTTGG | Amplification of *med16Δ::KAN-MX* |
| *MED16* amp RP | GAGAGAAATGCGTACCCTTGG | |
| *MED16* conf FP | CCAATGTTGACAGAACTGGC | Confirmation of *med16Δ::KAN-MX* insertion |
| *MED16* conf RP | CACTATCTGCCACGTAGATCG | |
| *KAN-MX* RP | CTTCCCATACAATCGATAGATTGTCG | Confirmation of *KAN-MX* insertion |
| *MED15* Chimeric FP | AGCGTATCGTTTCGTATAGTGCCGTACTCAAAGATCAAGGATTAAAACGCATTTAGGTGACACTATAG | Amplification of *med16Δ::KAN-MX* cassette |
| *MED15* Chimeric RP | TGGTGACCATAACACCAAACGAAGTAACTTCAAAAGTATCAAAAGTATGGTAATACGACTCACTATAGGG | |
| *MED15* Screening FP | TGTACAGTCCACGGATGGTGCAGAAG | Confirmation of *med15Δ::KAN-MX* insertion |
| *KAN-MX* RP | GAGTAACCATGCATCATCAGGAGTACGG | |
| *MED15* FP -530 | GGAACTACACAGACTTCCTTGTCGCGATAC | |

| | | |
|---|---|---|
| *MED15* R4 +3278 | GGTGACCATAACACCAAACGAAGTAACTTC | Amplification of *med15-Δdomain:: URA3* |
| *MED15 F1 -68* | GCGTATCGTTTCGTATAGTGCCGTAC | Confirmation of *med15-Δkix:: URA3* |
| *MED15 R1 +400* | GCAACCTGTTGCCTTGCCTG | |
| *MED15* F2 +296 | CGTGGAACAGCACCATATTAACAAC | Confirmation of *med15-Δabd1:: URA3* |
| *MED15* R2+1186 | GCGGCAGCATTAGGTGTAG | |
| *MED15* F3 +726 | GCTAATAACAACAACAACGGCCTC | Confirmation of *med15-Δabd2:: URA3* |
| *MED15* R2+1186 | GCGGCAGCATTAGGTGTAG | |
| *MED15* F3 +726 | GCTAATAACAACAACAACGGCCTC | Confirmation of *med15-Δabd3:: URA3* |
| *MED15* R4 +3278 | GGTGACCATAACACCAAACGAAGTAACTTC | |
| *MED17* amp FP | CAGCCCTTTGAAAAAGTAGAACTGC | Amplification of *MED-17-9xMyc::TRP1* |
| *MED17* amp RP | CATAACACCGCAGCCTAAAATTGC | |
| *MED17* Conf FP1 | CATTCACATCAAGTTTTCAGCTGGTACG | Confirmation of *MED-17-9xMyc::TRP1* |
| *MED17* Conf RP1 | CGAAGATAAGATCTTGTCCTTTTGTACGTATGC | |
| *MED3* chimeric FP | CTCTAGACCTGAACAATCTGGAATTAGGTGGTCTGAACATGGATTTCTTGGGGGGAGGCGGGGGTGGA | Amplification of *MED3-3xFLAG::HIS3* cassette |
| *MED3* chimeric RP | AATAGAAGATTATACAGATAATTACTATCTTGGATACATAGATGCACCAGGAATTCGAGCTCGTTTAAAC | |
| *MED3* scng FP | GCATGAATAACATGAATAACGG | Confirmation of *MED3-3xFLAG::HIS3* |
| *MED3* scng RP | GTAAAATTAGTCATTCTTCCACG | |
| *MED15*-C-*mAID*-F | GTTCAGAACAATTCAATGTATGGGATTGGAATAATTGGACAAGTGCTACTCGTACGCTGCAGGTCGAC | Amplification of *MED15-mAID* cassette |
| *MED15*-C-*mAID*-R | ACCAAACGAAGTAACTTCAAAAGTATCAAAAGTATGGAAACTTCAAATGTTCGATGAATTCGAGCTCG | |
| *MED15* Conf F | AGTAGATTCTCCTGATGACC | Confirmation of *MED15-mAID* tagging |
| *MED15* Conf R | CTAGTACTGATGATAGTCAAGTCC | |

| | | |
|---|---|---|
| *MED16-mAID*-Cterm F | TGTATACAAGACTGTGCATATGTTCAGGT ATGCTTTTTGAGATGGACGGCCGTACGCT GCAGGTCGAC | Amplification of *MED16-mAID* cassette |
| *Med16-mAID*-C term R | GTGAAATGTTTAAAACAATTCTATACAAAA CTATGCTATAGTACTAATAATCGATGAATT CGAGCTCG | |
| *MED16* Conf F | CTGAGTGTATCAGAAATATCC | Confirmation of *MED16-mAID* tagging |
| *MED16* Conf R | CAAAGGAACATTATTGCTC | |
| *RPB3* C-term F | TATTGTTAGCTCTGACACAGATG | Amplification of *RPB3-mCherry* cassette |
| *RPB3* C-term R | GTAGATTTGACATTCGGTAGTTC | |
| *RPB3* Conf F: | GATCAAGTTGTCGTCAGAGGTATCG | Confirmation of *RPB3-mCherry* tagging |
| *RPB3* Conf R: | ATTAGTAGACGAACTAAGTCACG | |
| *HSF1* Cterm 1 F | TGACCACAGTTATTCCACC | Amplification of *HSF1-mNeonGreen* cassette |
| *HSF1* Cterm 1 R | GCAGTTCAACCTCACTCG | |
| *HSF1*_Conf 2 _F | ACGACAATAACACTAGTGAGG | Confirmation of Hsf1-mNG tagging |
| *HSF1*_Conf 2_R | CTCAGGCTCTCACTAGCTC | |
| **Primers used for *med2-∆IDR*, *med3-∆IDR* strain construction by CRISPR** | | |
| **sgRNA Name** | **Sequence (5' → 3')** | **Purpose** |
| MED2 gAssembly 1 FP | gactttAaagaataagaacaacaacg | *MED2* sgRNA 1 |
| MED2 gAssembly 1 RP | aaaccgttgttgttcttattcttTaa | |
| MED2 gAssembly 2 FP | gactttaacaacaacaacaacaacag | *MED2* sgRNA 2 |
| MED2 gAssembly 2 RP | aaacctgttgttgttgttgttgttaa | |
| MED3 gAssembly 1 FP | gactttCCCATATTCATATTGtactg | *MED3* sgRNA 1 |
| MED3 gAssembly 1 RP | aaaccagtaCAATATGAATATGGGaa | |
| MED3 gAssembly 2 FP | gactttGTTTAATTGTGACTGTAGCG | *MED3* sgRNA 2 |
| MED3 gAssembly 2 RP | aaacCGCTACAGTCACAATTAAACaa | |
| **For *MED2, MED3* Donor Template amplification** | | |
| **Name** | **Sequence (5' → 3')** | |
| *MED2* donor template | CGAAACCAAAGGCTTTGACGATAATGACT CAGGTAACAACTACAATGACATAAATATTT | *MED2* IDR (290-400 aas) deletion |

| | | |
|---|---|---|
| | CTTCTATTGAAAACAACATAAATAATAATA TCAATAGCACCGAATACTTGACATTAAATG ATTTCAACGACCTTAATATTGACTGGTCGA CCACTGGAGATAATGGCGAATTAGACCTC AGCGGCTTCAATATATAGCTTG | |
| *MED2 FP* | CGAAACCAAAGGCTTTGACG | For *MED2* template amplification |
| *MED2* RP | CAAGCTATATATTGAAGCCGCTG | |
| *MED3* donor template | GCCCACAAATGTTATGGGCACGCCGCTG ACCAACATGATGTCTCCCATGGGGAACGC ATACTCAATGGGAGCTCAAAACCAAGGGG GACAAGTATCTATGGGGGGGAAAGAACT GGATTCTCTAGACCTGAACAATCTGGAAT TAGGTGGTCTGAACATGGATTTCTTGTGA CTGGTGCATCTATGTATCCAAGATAGTA | *MED3* IDR (301-374 aas) deletion |
| *MED3* FP | GCCCACAAATGTTATGGG | For *MED3* template amplification |
| *MED3* RP | TACTATCTTGGATACATAGATGC | |
| *MED2 IDR* scng FP | GATCCTCCCAAATAAACTGC | For confirmation of *med2-ΔIDR* |
| *MED2 IDR* scng RP | GCTGTTGTTGTTGTTCTCG | |
| *MED3 IDR* scng FP | CAACCCAACTCCTAGCATGA | For confirmation of *med3-ΔIDR* |
| *MED3 IDR* scng RP | TCCAGATTGTTCAGGTCTAG | |
| **For *MED2 or MED3* tagging with 9xMyc** | | |
| *MED2* Chimeric FP | GGTCGACCACTGGAGATAATGGCGAATTA GACCTCAGCGGCTTCAATATAtccggttctgctg ctag | For amplification of *MED2-9xMYC::TRP1* cassette |
| *MED2* Chimeric RP | GAATGCACAACACGGTTTACAAGTCAATA GTTAACAATAGGAAGACCAAGcctcgaggcca gaagac | |
| *MED2* scng FP | CAATCCTGGAGATAACCCTCC | For *MED2-9xMYC::TRP1* screening |
| *MED2* scrng RP | CATGCATCTCTCACATGACG | |
| *MED3* Chimeric FP | CTCTAGACCTGAACAATCTGGAATTAGGT GGTCTGAACATGGATTTCTTGtccggttctgctg ctag | For amplification of *MED3-9xMYC::TRP*1 cassette |
| *MED3* Chimeric RP | AATAGAAGATTATACAGATAATTACTATCT TGGATACATAGATGCACCAGcctcgaggccag aagac | |

| | | |
|---|---|---|
| *MED3* scng FP | GAGCATGAATAACGATTTCCAGC | For *MED3-9XMYC::TRP*1 screening |
| *MED3* scrng RP | GAGCAATCGATGTTTACAGTCC | |

**Table 4. Primers Used for ChIP**

| Name | Sequence (5' → 3') |
|---|---|
| *ARS504 F* | GTCAGACCTGTTCCTTTAAGAGG |
| *ARS504 R* | CATACCCTCGGGTCAAACAC |
| *HSP104 UAS F -266* | CTTAAACGTTCCATAAGGGGC |
| *HSP104 UAS R -195* | TGCAGTTCTTTGAGATGGGCC |
| *HSP104 Prom F -130* | GCATTGTAATCTTGCCTCAATTCC |
| *HSP104 Prom R -70* | GTTATTGCTGATTCGATTCAAGG |
| *HSP104 ORF F +1469* | CCCTTGATGCTGAACGTAGATATG |
| *HSP104 ORF R +1621* | CCACATTTTGGATCATGGAGTTG |
| *HSP104 3'UTR F +2676* | AGGTGATGACGATAATGAGGACAG |
| *HSP104 3'UTR R +2839* | TCTTTTGCTCGGGTGTCAAGTTC |
| *HSP82 Prom F -157* | TCCGCCACCCCCTAAAAC |
| *HSP82 Prom R -113* | TGAGGAGGTCACAGATGTTAAGAATT |
| *HSP82 ORF F +1392* | GCCAGAACACCAAAAGAACATCTAC |
| *HSP82 ORF R+1522* | ATTCATCAATTGGGTCGGTCAAG |
| *HSP82 3'UTR F +2036* | ATGAGGATGAAGAAACAGAGACTGC |
| *HSP82 3'UTR R+2297* | ACACACTAGACGCGTCGGAATAG |
| *SSA4 UAS F -374* | GCCGCACATCCATTCCGGTATG |
| *SSA4 UAS R -291* | CGGGCAAAAGATATCCGCTTTG |
| *SSA4 Prom F -246* | AGTTCCTAGAACCTTATGGAAGCAC |
| *SSA4 Prom R +35* | GTTGTACCTAAATCAATACCAACAGC |
| *SSA4 ORF F +816* | GTCTTCGTCTGCTCAGACATC |
| *SSA4 ORF R +946* | CCACTGGCTCCAATGTAGATC |
| *SSA4 3'UTR F +1762* | GAGGAATACAAGGAAAGGCAAAAG |
| *SSA4 3'UTR R +2079* | TTAAACTCTGGCTTATGACGATGAG |

**Table 5. Primers Used for Taq I - 3C**

| Name | Sequence (5' → 3') |
|---|---|
| *ARS504 F* | GTCAGACCTGTTCCTTTAAGAGG |
| *ARS504 R* | CATACCCTCGGGTCAAACAC |
| *HSP12 F-47* | ACGTATAAATAGGACGGTGAATTGC |
| *HSP12 R-47* | TTCAGAAGCTTTTTCACCGAATC |
| *HSP82 F+740* | AATTAGTCGTCACCAAGGAAGTTG |
| *HSP82 R+740* | AATGCTTAACGTACAATGGGTCTTC |
| *HSP82 F+2189* | ATGAGGATGAAGAAACAGAGACTGC |
| *HSP82 R+2189* | ACACACTAGACGCGTCGGAATAG |
| *HSP104 F-63* | AGGCATTGTAATCTTGCCTCAATTC |
| *HSP104 R-63* | ATCGTTAGAGCCCTTTCTGTAAATTG |
| *HSP104 F+782* | GTAAGACCGCTATTATTGAAGGTG |
| *HSP104 R+782* | TTCTTCGATTTCCTTCAAAACACC |
| *HSP104 F+1550* | CCCTTGATGCTGAACGTAGATATG |
| *HSP104 R+1550* | CCACATTTTGGATCATGGAGTTG |
| *HSP104 F+2756* | AGGTGATGACGATAATGAGGACAG |
| *HSP104 R+2756* | TCTTTTGCTCGGGTGTCAAGTTC |
| *HSP26+221 F* | TATGATCCCAGAGATGAAACC |
| *HSP26 +221 R* | GAAACCGAAACCAGATGG |
| *SSA2 F+1368* | TCTCTACTTATGCTGACAACCAACC |
| *SSA2 R+1368* | TTCAATTTGTGGGACACCTCTTG |
| *SSA4 F-268* | ACACGAAAGATATCTCAACTCTAGCC |
| *SSA4 R-268* | TGTTACTGTCGTCAAACTAAGGAG |
| *SSA4 F+198* | GCCTTCTTATGTGGCTTTTACTGAC |
| *SSA4 R+198* | TTTACGTCCGATCAGACGCTTAG |
| *SSA4 F+2255* | ATAAGAAAGTCATCGCCAAACAAC |
| *SSA4 R+2255* | GTGTTAAACTCCGGTCAAAAGAAAC |
| *UBI4 F+524* | GTAAGCAGCTAGAAGATGGTAGAACC |
| *UBI4 R+524* | TGAATTTTCGACTTAACGTTGTCG |

**Table 6. Primers Used for RT-qPCR**

| Name | Sequence (5' → 3') |
|---|---|
| *HSP104 ORF F+1646* | CAGCTGCAAGATTGACTGGTATCC |
| *HSP104 ORF R+1799* | CCTGATCTAGACAATCTAACGGC |
| *HSP82 ORF F+290* | CAAGTCTGGTACCAAAGC |
| *HSP82 ORF R+453* | CAGTGAAAGAACCACCAGC |
| *SSA4 ORF F+815* | GTCTTCGTCTGCTCAGACATC |
| *SSA4 ORF R+946* | CCACTGGCTCCAATGTAGATC |
| *SCR1 F+385* | CGGCCGGGATAGCACATATC |
| *SCR1 R+438* | CGCCGAAGCGATCAACTTG |
| *BTN2 ORF F+555* | GTTTTTGTTATTGGCTGTGGAG |
| *BTN2 ORF R+649* | CTTCCTCATGCTTAACTAAACC |
